# A Machine Learning-Based Investigation of Integrin Expression Patterns in Cancer and Metastasis

**DOI:** 10.1101/2024.09.19.613933

**Authors:** Hossain Shadman, Saghar Gomrok, Qianyi Cheng, Yu Jiang, Xiaohua Huang, Jesse D. Ziebarth, Yongmei Wang

## Abstract

**Background:** Integrins, a family of transmembrane receptor proteins, play complex roles in cancer development and metastasis. These roles could be better delineated through machine learning of transcriptomic data to reveal relationships between integrin expression patterns and cancer.

**Methods:** We collected publicly available RNA-Seq integrin expression from 8 healthy tissues and their corresponding tumors, along with data from metastatic breast cancer. We then used machine learning methods, including t-SNE visualization and Random Forest classification, to investigate changes in integrin expression patterns.

**Results:** Integrin expression varied across tissues and cancers, and between healthy and cancer samples from the same tissue, enabling the creation of models that classify samples by tissue or disease status. The integrins whose expression was important to these classifiers were identified. For example, ITGA7 was key to classification of breast samples by disease status. Analysis in breast tissue revealed that cancer rewires co-expression for most integrins, but the co-expression relationships of some integrins remain unchanged in healthy and cancer samples. Integrin expression in primary breast tumors differed from their metastases, with liver metastasis notably having reduced expression.

**Conclusions:** Integrin expression patterns vary widely across tissues and are greatly impacted by cancer. Machine learning of these patterns can effectively distinguish samples by tissue or disease status.

## Background

Integrins are a family of transmembrane receptor proteins that facilitate adhesion and regulate essential cell activities such as cell communication, migration, survival, and proliferation.^1–3^ Structurally, integrins are heterodimers composed of an α and a β subunit.^3^ In humans, there are 18 known α subunits and 8 known β subunits that form at least 24 functional heterodimers. They have traditionally been subdivided into 5 distinct classes based on ligand interaction and other features, such as the structure of the α subunit (Supplementary Fig. 1).^4,5^

Due to the importance of integrins in cell communication mechanisms and how the use of these mechanisms by tumor cells evolves during disease progression, it is not surprising that integrins have been widely implicated as key players in cancer development.^6–10^ In breast cancer, for example, links between integrins and tumor development have been established through studies showing that deletion of integrins *β1* and *β4*, respectively, inhibits tumor initiation and progression to invasive adenocarcinoma.^11^ The metabolic changes associated with breast cancer and metastasis also involve integrin signaling, including hypoxia-inducible factors targeting integrins and modifying their signaling pathways during hypoxia.^12^ Integrins have been implicated in organotropism of breast cancer metastasis, as the specific integrins expressed in exosomes released by tumors have been associated with preparing the premetastatic niche in different tissues.^13^

Despite the well-established connection between integrins and cancer, the specifics of relationships between cancer and the expression of individual integrins are complex, with single integrins seeming to have opposite impacts in different cancers and different integrins having various functions during a single cancer type’s progression.^6^ For example, ITGA3 has been identified as both a risk factor and a protective factor in different cancers, as high expression of ITGA3 was linked to poor prognosis in some cancers and better prognosis in others.^14,15^ Another example of the complex behavior of integrins in cancer is their expression in breast cancer.^9,16^ Some integrins, such as *α11*, *β1*, and *β3*, were found to promote tumor cell invasion, progression, and metastasis in breast cancer,^10,17–19^ while others, such as *α7*, are down regulated in breast cancer.^9^ Because of this complexity, a systematic investigation of how integrin expression patterns are associated with cancer development, progression, and metastasis would be valuable.

The advent of next-generation sequencing technologies, such as RNA-Seq, has provided enormous amounts of genetic and transcriptomic data that can be leveraged to increase understanding of the roles of different classes of genes in cancer. In addition to the general increase in the understanding of gene expression offered by this data, quantification of mRNA expression has the potential to improve cancer diagnosis and treatment specifically, as mRNA expression alone is the basis for several current clinical tools including PAM50,^20–22^ Oncotype DX,^23^ and MammaPrint.^24^ Next-generation sequencing has also led to the introduction of several large-scale projects including the Genotype Tissue Expression (GTEx)^25^ Project, The Cancer Genome Atlas Project (TCGA)^26^, and AURORA^27^, a study of metastatic breast cancer, that aim to systematically characterize gene expression across a variety of tissues and conditions. The data generated by these projects provides an opportunity for data mining to investigate the changes in gene expression that underlie cancer development and metastasis.

Here, we report such a study as we use these data sets to comprehensively study integrin expression patterns and examine how integrin expression varies across healthy tissues, primary tumors, and metastasis. Our analysis is divided into two main sections. In the first section, we compare integrin expression across a set of 8 healthy tissues and their corresponding tumors. In the second section, we focus on breast cancer and compare integrin expression in healthy breast tissue, primary breast tumors, and breast cancer metastases.

## Materials and Methods

### Data Sources

Most analysis in this work focused on expression of 27 integrin genes: 18 α subunits, 8 β subunits, and the integrin-like gene ITGBL1. Except for data from AURORA,^27^ this study analyzed RNA-Seq data normalized with the TOIL^28^ Recompute pipeline. Specifically, we used the “gene expression RNAseq RSEM tpm” file of the TCGA-TARGET-GTEx cohort downloaded from XenaBrowser.^29,30^ GTEx is our main source of healthy tissue data and TCGA is a source of cancer tissue data. We selected tissues to analyze by choosing all non-brain organs that had sample size > 100 in both GTEx and TCGA projects, resulting in the selection of 8 tissues: breast, colon, liver, lung, pancreas, prostate, stomach, and testis. Supplementary Table 1 summarizes the samples sizes of each healthy tissue and cancer studied.

Metastatic breast cancer data from the AURORA US Metastasis Project^27^ was downloaded as the AURORA Upper Quantile Normalized (UQN) RNA-Seq dataset from the GEO repository (GSE209998)^31,32^ in February 2023. We transformed the gene expression values by computing log_2_(x+1) where x is the value in the downloaded file. This dataset consists of 44 primary tumor and 79 metastatic samples from 53 patients. Metastasis samples came from 19 locations, with liver (n = 18), lymph node (n = 11), brain (n = 9), and lung (n = 8) being the most common.

### Machine Learning and Statistical Methods t-SNE visualization

T-distributed stochastic neighbor embedding (t-SNE) was used for dimension reduction of the expression of the 27 integrins using the Python scikit-learn^33^ library with perplexity parameters between 40 and 50. The first two dimensions in the t-SNE space were used for visualization.

### Random Forest classification

We employed Random Forest (RF) models in several ways: (i) to classify samples by tissue or cancer type using multiclass or one-vs-all classifiers, (ii) to classify samples from the same tissue as healthy or cancer, and (iii) to classify samples as primary breast tumor or metastasis. RF models were implemented with the Python scikit-learn^33^ Random Forest classifier and used a 50%/50% training/test split. To address class imbalance, we set the class_weight parameter to ‘balanced’. This setting yielded results consistent with those obtained when employing over- or under-sampling techniques using the imbalance-learn library. We conducted 500 iterations for each RF model that each had a randomly selected training/test split. Model validation metrics, such as accuracy and area under the ROC curve (AUROC), and feature importances are averaged over the 500 iterations.

### Co-expression analysis

For gene co-expression analysis, we computed Pearson correlation coefficients (R) between the expression of genes across samples. Integrin-integrin pairs were considered co-expressed if the R between their expression was greater than +0.6. The mean Pearson coefficient between every integrin-integrin pair in the GTEx breast and TCGA BRCA primary tumor datasets were 0.14 and 0.24, respectively. For integrin-integrin pairs, we searched STRING-db^34^ to check for known associations in human. Graphviz^35^ was used for co-expression network visualization. We also examined co-expression relationships between integrins and ∼19000 protein coding genes that were selected using BioMart.^36^

### Statistical tests

The expression of each integrin was categorized as under- or over-expressed in cancer based on whether the expression of the integrin was lower in tumor samples compared to healthy/normal samples, or vice versa. For all plots associated with RF models, an asterisk denotes that the expression difference was significant, where significance was defined as the Bonferroni adjusted p-value of a t-test (ttest_ind from scipy.stats and statsmodels.test.multi.multipletests) falling below 0.05. Similar analysis was performed for metastatic breast cancer data, with under-expression defined as having lower expression in metastatic samples. One-way ANOVA (f_oneway from scipy.stats) followed by Tukey post-hoc tests (pairwise_tukeyhsd from statsmodels.stats.multicomp) were used to identify integrins with significantly different expression in primary breast tumors and four metastasis locations (liver, lymph node, brain, and lung).

## Results

### Integrin expression patterns can differentiate tissue compartments

Unperturbed human tissues differ markedly in the composition of their extracellular matrices, and this could be expected to correspond to variation in integrin expression. To explore this notion, we calculated mean mRNA expression levels of integrins in 8 selected healthy tissues from the GTEx database (Fig. 1a). There was variation in both the overall mean expression of the different integrins and the expression of each integrin across tissues. To further explore the relationship between integrin expression and tissue, we created a t-SNE plot of the samples in the 8 tissues using only integrin expression as input features (Fig. 1b). The t-SNE plot shows that, in general, samples from each tissue formed clusters that were relatively distinct from those of other tissues, indicating that integrin expression patterns are tissue specific.

**Fig. 1:**
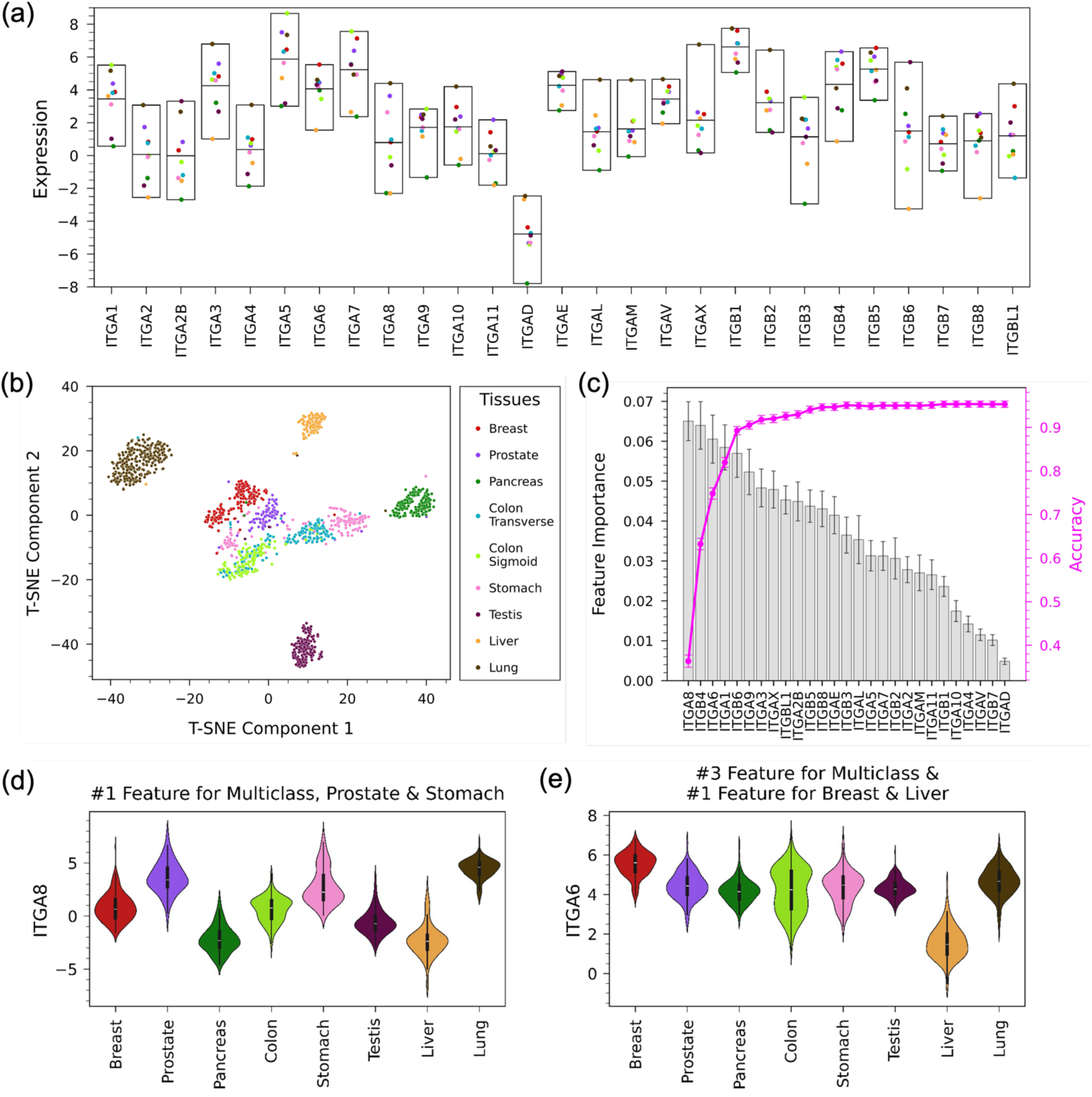
Variation of integrin expression across healthy tissues. (a) Mean expression of integrins in 8 healthy tissues in the GTEx dataset. The middle line of each box shows the mean of means. (b) t-SNE plot based on integrin expression in healthy tissues shows clustering of samples based on tissue. (c) Feature importance of integrins in a multiclass Random Forest model using a 50%/50% training/test split that classifies healthy samples by tissue. The magenta line shows the change in the accuracy of the model as features are added one-by-one from the most important feature to the least. The accuracy of the model using all 27 integrins is 0.953 ± 0.007 and AUROC is 0.995 ± 0.002. (d) Violin plots of expression across tissues for ITGA8, the top feature in the multiclass RF model and in one-vs-all RF models for prostate and stomach. (e) Violin plots of expression across tissues for ITGA6, the third-ranked feature in the multiclass RF model and the top feature in the RF models for breast and liver.

We then applied a multiclass Random Forest (RF) model to classify GTEx samples according to tissue based on integrin expression. This model had a high accuracy (0.953 ± 0.007) and AUROC (0.995 ± 0.002), suggesting that integrin expression patterns are unique to tissue compartments. The confusion matrix of the model’s predictions showed that performance was strong across tissues, as classification accuracy was greater than 95% for 6 of the 8 tissues (Supplementary Fig. 2). Accuracy for prostate (84%) and stomach (88%) was somewhat lower, as the model incorrectly classified ∼7% of the samples of these tissues as colon. In addition to making predictions, random forest models can be used to determine how features contribute to the model’s predictions. Specifically, we used the feature importance of each integrin in the multiclass RF model to determine the integrins that were most important when classifying samples by tissue (Fig. 1c). Integrins with high feature importance tended to have particularly high or low expression in one or two tissues (Fig. 1d, e and Supplementary Fig. 3), with ITGA8, for example, having relatively high expression in lung and prostate and low expression in liver and pancreas (Fig. 1d). The maximum feature importance in the multiclass RF model was relatively low (0.07), indicating that the model was not highly dependent on the expression of a single integrin (Fig. 1c).

We also created one-vs-all RF models for each of the 8 tissues to further investigate the ability of integrin expression patterns to classify samples by tissue type and identify the integrins that were important in distinguishing individual tissues (Table 1). Like the multiclass model, the one-vs-all RF models had high accuracy (> 95% for all 8 tissues). However, the feature importances of the one-vs-all models showed different behavior than the multiclass model, as many one-vs-all models were reliant on the expression of one or two integrins (Supplementary Fig. 4). In the one-vs-all model for stomach, for example, the feature importance of ITGA8 was 0.175, a value almost three times that of any other integrin (Supplementary Fig. 4g). While the RF models described to this point used the expression of all integrins as input features, models with high accuracy could be achieved using only a handful of integrins. Fig. 1c shows how the accuracy of the multiclass RF model changes as the number of features is increased (i.e., integrins are added to the model one at a time in order of their feature importance in the multiclass model with all integrins). Achieving an accuracy of 90% for the multiclass model required including the expression of only 5 integrins, and the accuracy reached a plateau after ∼10 integrins are added to the model. For most one-vs-all models, the expression of only 1 or 2 integrins was needed for an accuracy higher than 0.9 (Supplementary Fig. 4).

**Table 1:**
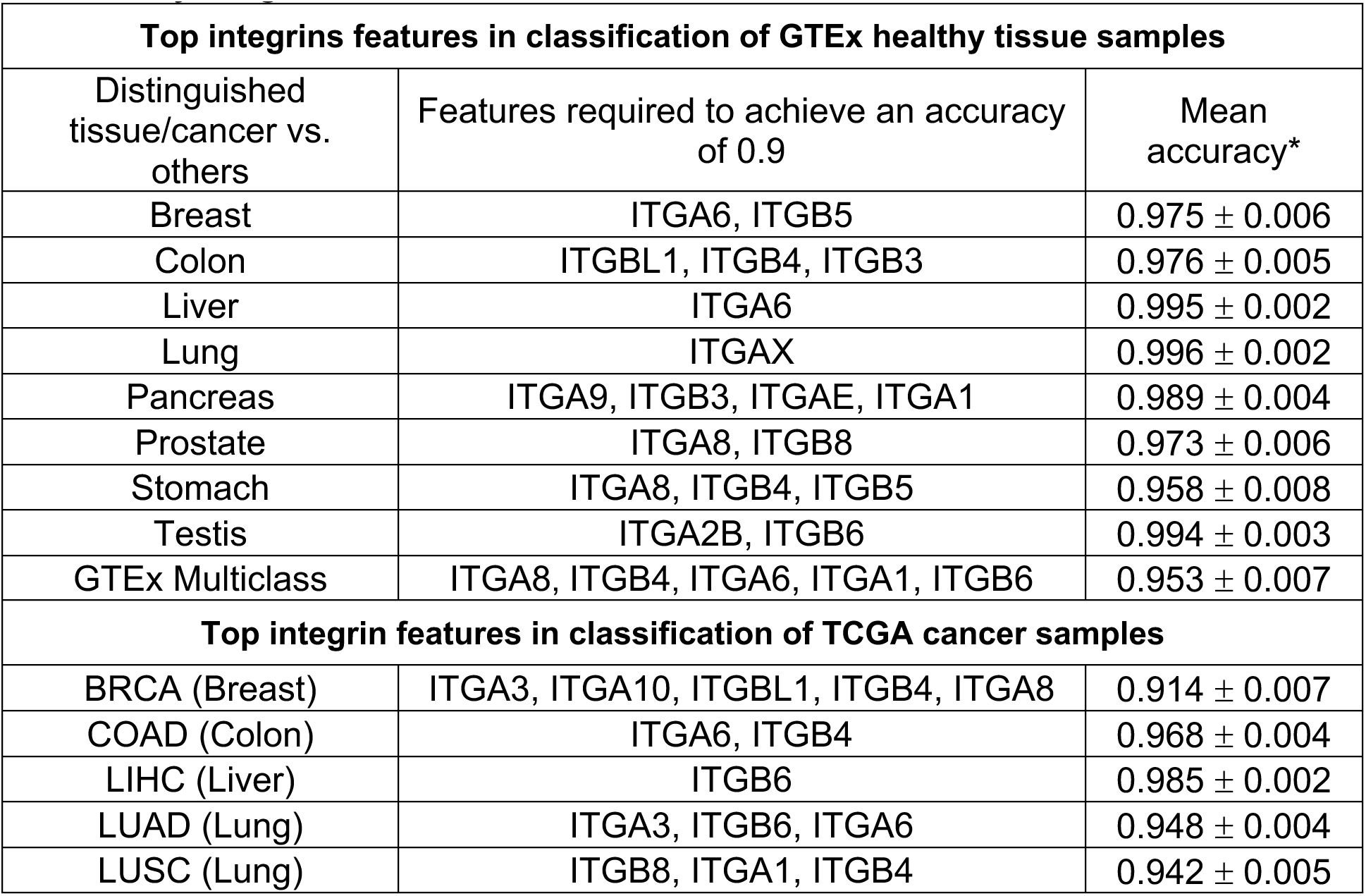
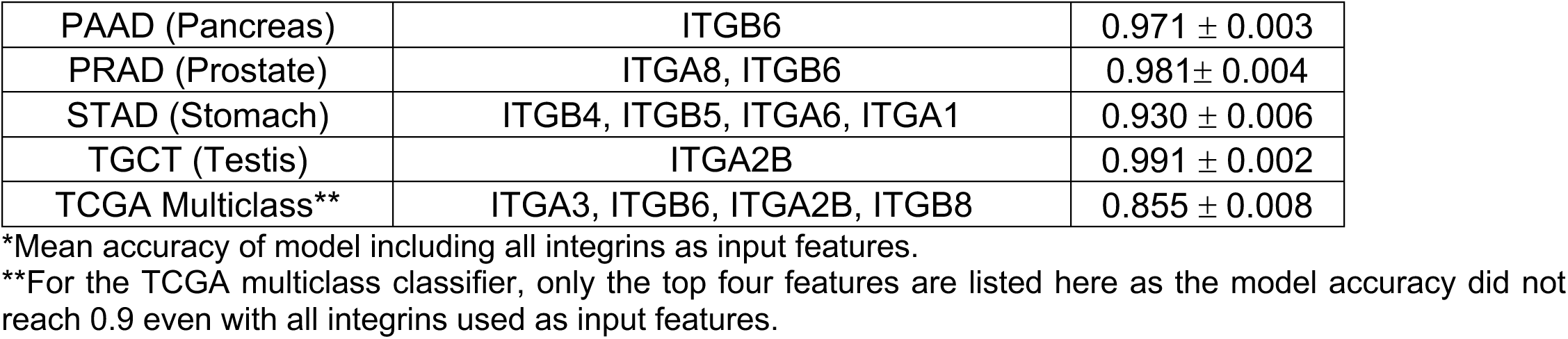
Key integrins in Random Forest classification models.

| **Top integrins features in classification of GTEx healthy tissue samples** |  |  |
| --- | --- | --- |
| Distinguished tissue/cancer vs. others | Features required to achieve an accuracy of 0.9 | Mean accuracy* |
| Breast | ITGA6, ITGB5 | 0.975 ± 0.006 |
| Colon | ITGBL1, ITGB4, ITGB3 | 0.976 ± 0.005 |
| Liver | ITGA6 | 0.995 ± 0.002 |
| Lung | ITGAX | 0.996 ± 0.002 |
| Pancreas | ITGA9, ITGB3, ITGAE, ITGA1 | 0.989 ± 0.004 |
| Prostate | ITGA8, ITGB8 | 0.973 ± 0.006 |
| Stomach | ITGA8, ITGB4, ITGB5 | 0.958 ± 0.008 |
| Testis | ITGA2B, ITGB6 | 0.994 ± 0.003 |
| GTEx Multiclass | ITGA8, ITGB4, ITGA6, ITGA1, ITGB6 | 0.953 ± 0.007 |
| **Top integrin features in classification of TCGA cancer samples** |  |  |
| BRCA (Breast) | ITGA3, ITGA10, ITGBL1, ITGB4, ITGA8 | 0.914 ± 0.007 |
| COAD (Colon) | ITGA6, ITGB4 | 0.968 ± 0.004 |
| LIHC (Liver) | ITGB6 | 0.985 ± 0.002 |
| LUAD (Lung) | ITGA3, ITGB6, ITGA6 | 0.948 ± 0.004 |
| LUSC (Lung) | ITGB8, ITGA1, ITGB4 | 0.942 ± 0.005 |
| PAAD (Pancreas) | ITGB6 | $0.971 \pm 0.003$ |
| PRAD (Prostate) | ITGA8, ITGB6 | $0.981 \pm 0.004$ |
| STAD (Stomach) | ITGB4, ITGB5, ITGA6, ITGA1 | $0.930 \pm 0.006$ |
| TGCT (Testis) | ITGA2B | $0.991 \pm 0.002$ |
| TCGA Multiclass** | ITGA3, ITGB6, ITGA2B, ITGB8 | $0.855 \pm 0.008$ |

\*Mean accuracy of model including all integrins as input features.
\*\*For the TCGA multiclass classifier, only the top four features are listed here as the model accuracy did not reach 0.9 even with all integrins used as input features.

### Integrin expression patterns can differentiate solid tumor types

We then followed a similar procedure to examine integrin expression in TCGA primary tumor samples derived from the same tissue types analyzed earlier. Expression varied across integrins and across tumor tissues (Fig. 2a), like in the healthy samples. The t-SNE plot for 9 tumor types (there are two types of lung cancers: lung squamous cell carcinoma and lung adenocarcinoma) shows that the samples are relatively well separated based on their tissue lineage (Fig. 2b). However, they appear to be less clearly delineated than in the healthy tissues (Fig. 1b). This observation was confirmed by a multiclass RF model trained/tested on the tumor samples, which had an accuracy of 0.86 ± 0.01, below the 0.95 achieved for healthy samples (Fig. 2c). The lower accuracy for the tumor samples can be mostly explained by the model tending to incorrectly classify samples as breast cancer (Supplementary Fig. 5). One-vs-all classifiers were able to predict the origin of tumor samples with accuracy close to that of healthy samples for many tissues (Table 1). In agreement with the multiclass RF confusion matrix, breast cancer had the lowest accuracy among the one-vs-all models. In general, the feature importance of the integrins in the tumor RF models (Fig. 2d, e; Supplementary Figs. 6 and 7) indicated similar behavior to that of the healthy samples (e.g., no integrins had high feature importance in multiclass model), although the specific integrins with high feature importance were different in many cases.

**Fig. 2:**
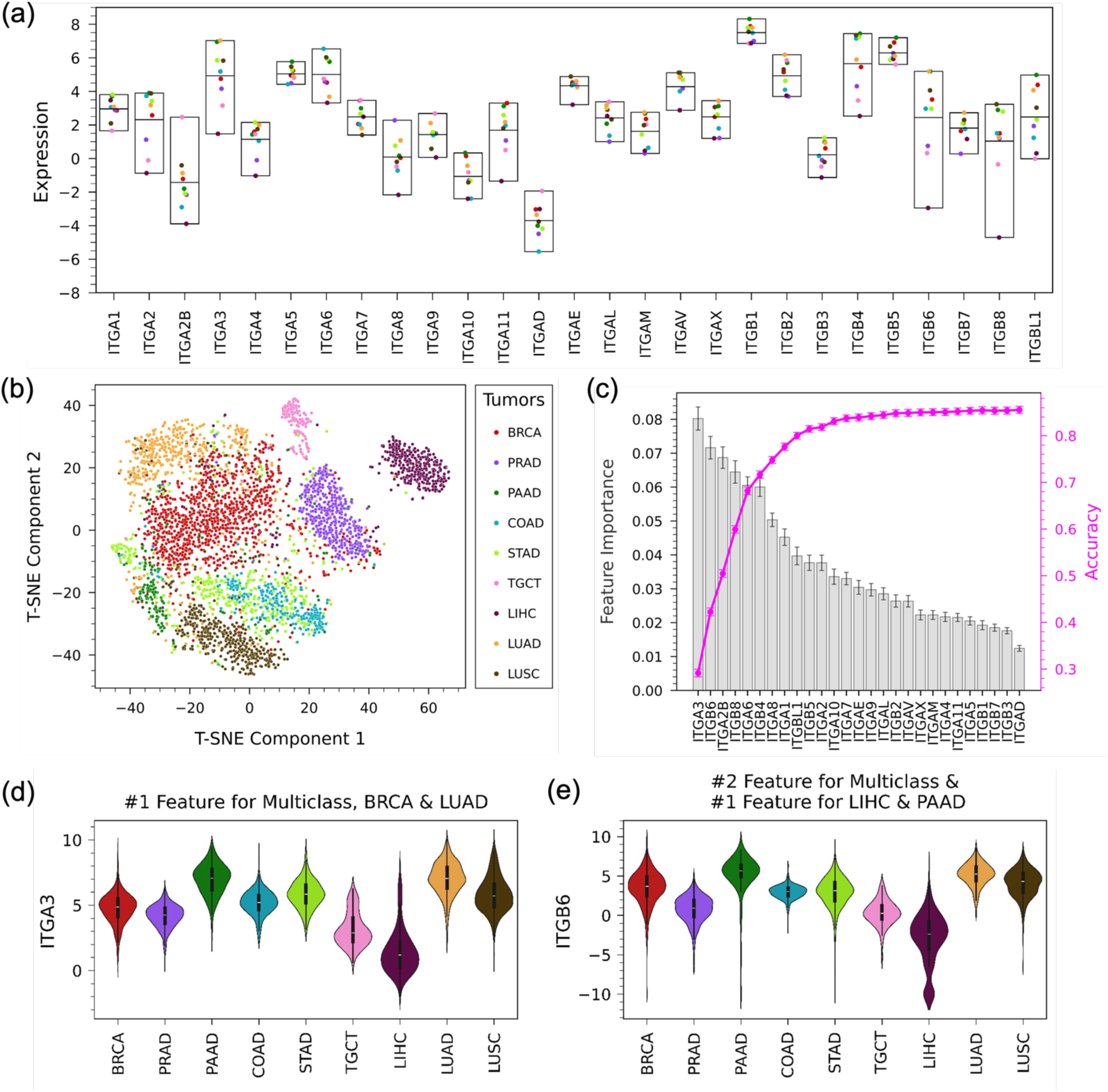
Variation of integrin expression across cancer types. (a) Mean expression of integrins in 9 cancer types in the TCGA dataset. The middle line of each box shows the mean of means. (b) t-SNE plot based on integrin expression in cancer shows clustering of samples based on cancer type, (c) Feature importance of integrins in a multiclass Random Forest model with a 50%/50% training/test split that classifies primary tumor samples by cancer type. The magenta line shows the model accuracy as features are added one-by-one from the most important feature to the least. The accuracy of the model using all 27 integrins is 0.855 ± 0.008 and AUROC is 0.982 ± 0.002. (d) Violin plot of expression across cancers of ITGA3, the top feature of the multiclass RF model and the one-vs-all RF models for BRCA and LUAD. (e) Violin plot of expression across cancers for ITGB6, the second ranked feature of the multiclass RF model and the top feature of one-vs-all RF models for LIHC and PAAD.

Taken together, the results from this section indicate that integrin expression patterns can be used to classify samples by tissue for both healthy and tumor tissues. In some cases, the success of the Random Forest models is due to the same integrins when classifying both healthy tissues and the tumors derived from these tissues (Table 1). In the one-vs-all models for prostate (ITGA8) and testis (ITGA2B), the same integrin was identified as the most important feature for classifying both healthy and tumor samples. However, the specific integrins with the highest feature importance in corresponding models were usually different, as, for example, there was no overlap in the top five integrins in the multiclass RF models for healthy and tumor samples.

### Changes in Integrin expression patterns from healthy tissue to tumor samples

To gain more insight into how integrin expression patterns change from healthy tissues to tumors, we looked at the difference between the mean expression of each integrin in GTEx healthy tissue samples and corresponding TCGA primary tumor samples (Fig. 3). The two subtypes of healthy colon tissue and the two lung tumor types were combined in both cases. As expected, the relationship between the development of cancer and integrin expression is complex, with cancer resulting in expression changes that vary across integrins for a given tissue and across tissues for a given integrin in most cases. One exception to this is pancreas, in which all integrins had higher expression in cancer than in healthy tissue, a behavior that has been previously reported.^8,37–40^ There are also some integrins (i.e., ITGA7, ITGA10, and ITGA2B) whose expression decreased in tumors for all tissues except pancreas and integrins (i.e., ITGA2 and ITGA11) whose expression increased in tumors for most tissues. To compare the variation in integrin expression, we calculated the standard deviation in mean expression for each integrin across tissues for GTEx samples and across tumors for TCGA (Supplementary Fig. 8). This standard deviation was higher for GTEx samples than TCGA samples for 22 of the integrins. Therefore, there was reduced variation in integrin expression across cancers for most integrins in comparison with healthy tissue counterparts.

**Fig. 3:**
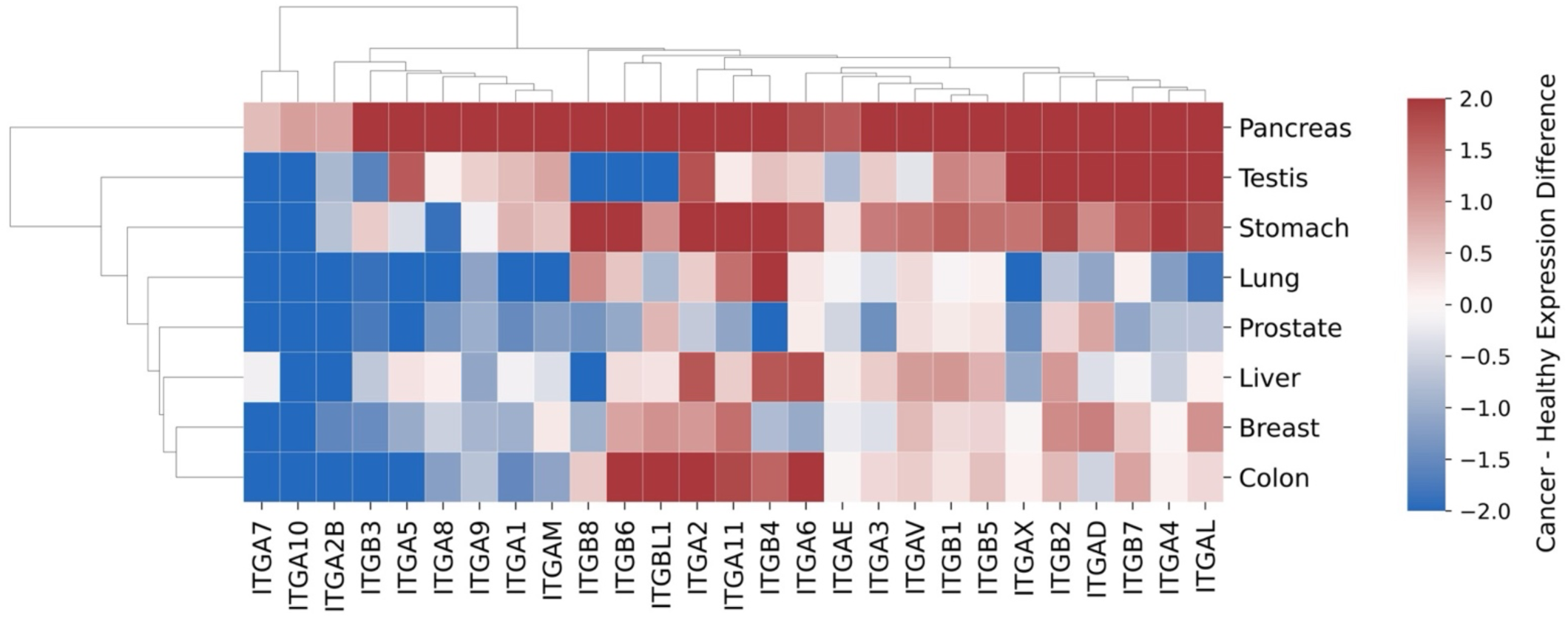
Impact of cancer development on integrin expression. The heatmap shows differences between the mean expression of an integrin in GTEx healthy tissue samples and the mean expression of that integrin in corresponding TCGA primary tumor samples. Red indicates higher expression in tumor samples, while blue indicates lower expression in tumor samples.

### Integrin expression differentiates healthy and cancer breast tissue samples

To further investigate the roles of integrins in cancer development, we focused our attention on breast cancer. We tested if integrin expression could classify healthy tissue from tumor samples. RF classifiers were trained to classify healthy versus tumors using two comparisons: (i) patient-matched tumor/normal tissue samples from TCGA BRCA (Fig. 4a) and (ii) TCGA BRCA primary tumor and GTEx healthy breast samples (Fig. 4b). In both cases, the RF classifier reached a high accuracy, around 0.96 for the patient-matched case and 0.99 for TCGA BRCA/GTEx. The top feature identified in both RF classifiers was ITGA7. ITGA7 and the other top features in these RF models tended to have lower expression in tumors than in healthy tissue in breast samples (Fig. 4c) and in several other organs (Fig. 3).

**Fig. 4:**
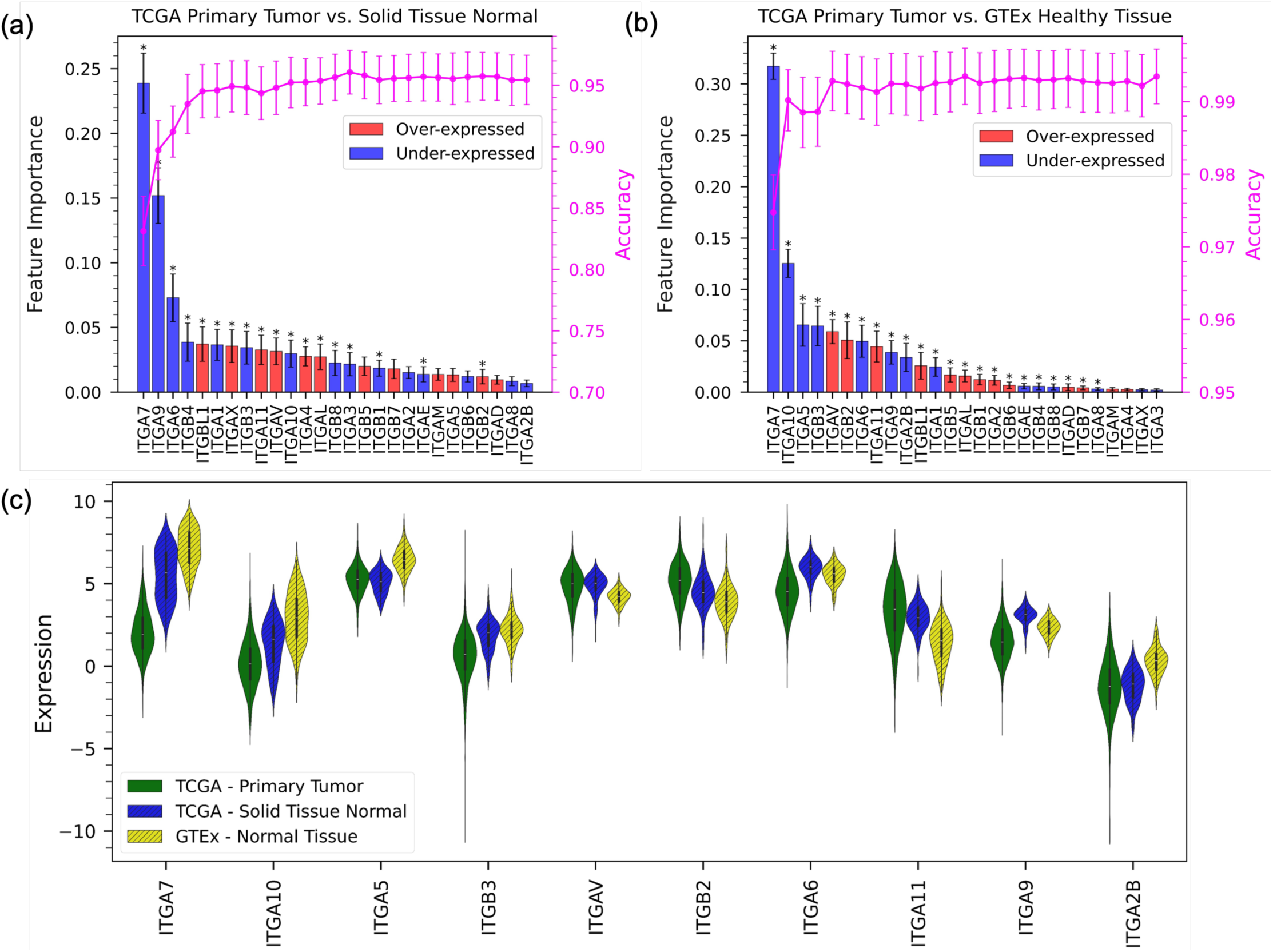
Integrin expression in healthy and cancer breast tissue. Feature importance and increase in accuracy with number of features for binary Random Forest models classifying samples as healthy or cancer for (a) primary tumor and solid tissue normal samples in the TCGA BRCA patient-matched data (accuracy of the model using all 27 integrins is 0.954 ± 0.020 and AUROC is 0.991 ± 0.006) and (b) TCGA BRCA primary tumor samples and GTEx healthy tissue samples (accuracy of the model using all 27 integrins is 0.993 ± 0.004 and AUROC is 0.999 ± 0.001). (c) Violin plots of the expression of integrins with high feature importance in the RF model for TCGA BRCA primary tumor and GTEx normal tissue samples.

Supplementary Fig. 9 presents similar results for RF classifiers of normal/tumor samples for the other tissue types investigated here (pancreas and testis were excluded due to having very few patient-matched samples). Breast tissue was unique among the tissues, as it was the only tissue with the same top feature in the RF classifiers trained using both the patient-matched TCGA and the TCGA/GTEx data. We investigated how the ability of ITGA7 to classify breast samples as healthy/cancer compared with two genes sets whose expression has been shown to be related to cancer development. Specifically, we created RF healthy/cancer breast sample classifiers using ITGA7 in combination with the PAM50 gene set^20–22^ (Supplementary Fig. 10a) or a set of genes from Donato *et al.*^41^ that are differentially expressed between hypoxic circulating tumor cell (CTC) clusters and normoxic CTCs (Supplementary Fig. 10b). In both classifiers, ITGA7 had the highest feature importance, ranking above all genes in both the PAM50 and Donato sets.

### Integrin co-expression in breast cancer

To further investigate changes in integrin expression in breast cancer, we examined co-expression relationships involving integrins in the GTEx breast and TCGA BRCA (primary tumor) datasets. Gene co-expression identifies genes with similar expression profiles that may indicate functional relationships or co-regulation.^42,43^ We studied co-expression between integrins and created networks of co-expressed integrin-integrin pairs in healthy (Fig. 5a) and cancerous (Fig. 5b) breast tissue. These co-expression networks were based solely on correlation in expression and did not include any previous biological knowledge, such as integrins that are known to form dimers. For both healthy tissues and cancers, the co-expression of integrins is common, with both healthy and cancer networks having over 20 co-expressed integrin-integrin pairs. Many of the co-expressed integrin-integrin pairs in both datasets have been previously linked in associations included in STRING-db^34^ (Supplementary Tables 2 and 3).

**Fig. 5:**
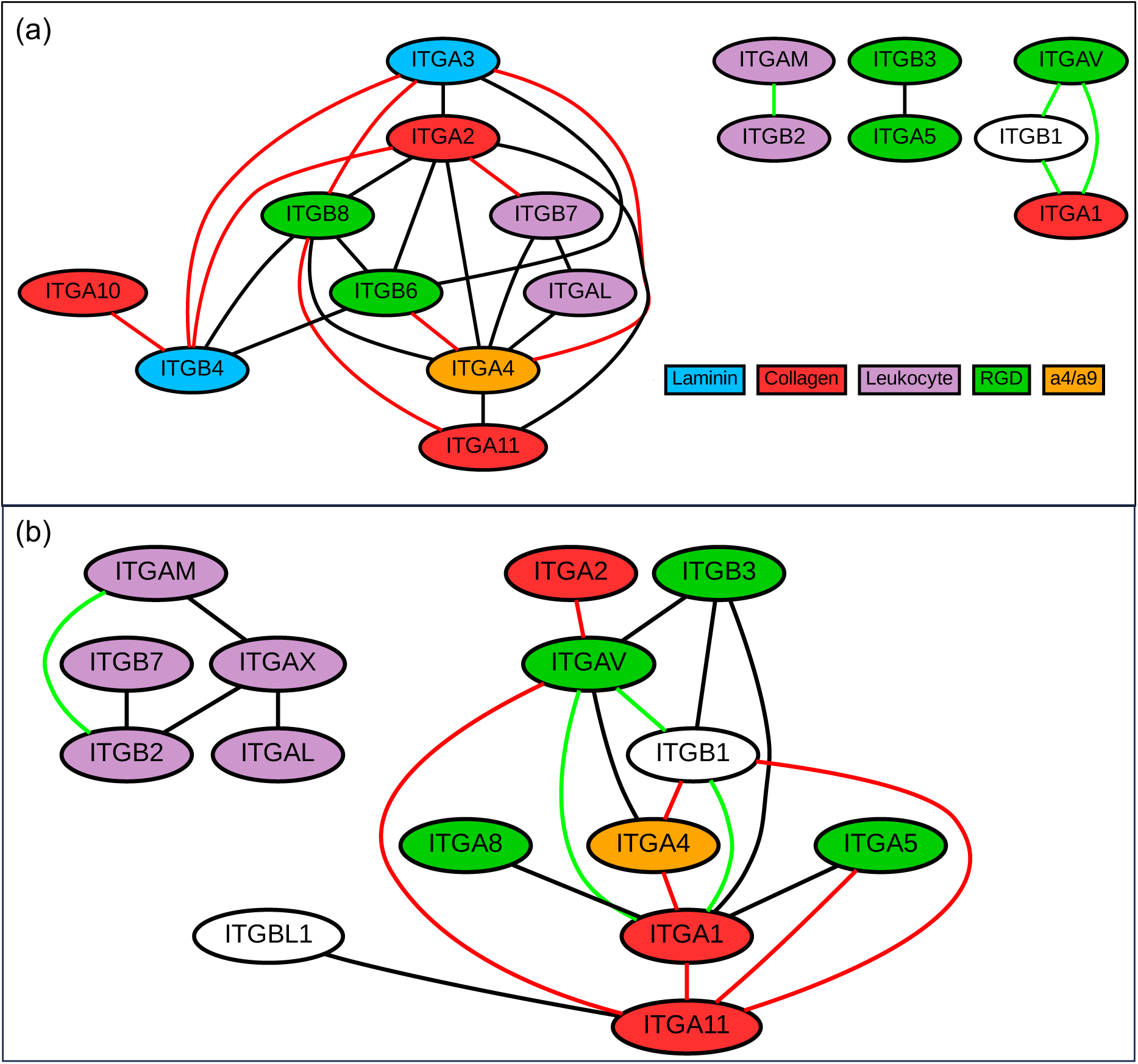
Integrin co-expression relationships in healthy breast tissue and breast cancer. Integrin-integrin co-expression networks in the (a) GTEx (breast) and (b) TCGA BRCA primary tumor datasets, with edges indicating pairs that have Pearson R ≥ 0.6 in the dataset. Green edges indicate integrin pairs that are co-expressed in both healthy and cancer datasets. Red edges indicate integrin pairs whose co-expression was strongly impacted by cancer (i.e., the difference between the R value in the healthy and cancer datasets was > 0.4).

Despite the general similarity in the structure of the co-expression networks for the two conditions, cancer development does greatly impact integrin-integrin co-expression. This impact can be observed through examination of the identities of the integrins in the co-expressed pairs, as ∼90% of the co-expressed integrin pairs are not conserved (i.e., Pearson R ≥ 0.6 in either healthy or cancer samples, but not both) across the two conditions. In many cases, the difference in the correlation coefficient of an integrin-integrin pair between healthy and cancer samples is large (integrin pairs connected by red edges in Fig. 5; Supplementary Tables 2 and 3). For example, ITGA3 and ITGB8 are highly co-expressed in healthy tissue (R = 0.72), but this relationship is absent in cancer (R = 0.00). Cancer does not disrupt the co-expression of all integrin-integrin pairs, as four co-expression relationships (ITGAM-ITGB2 and connections between ITGA1, ITGAV, and ITGB1, shown with green edges in Fig. 5) are conserved across the two networks (i.e., R ≥ 0.6 in both healthy and cancer samples). To examine if the impact of cancer on integrin co-expression was found more broadly, we investigated the co-expression between integrins and all protein coding genes (Supplementary Fig. 11). The impact of cancer on integrin-protein coding gene co-expression largely mirrored the behavior seen in the integrin-integrin co-expression networks. The co-expression with all protein-coding genes for integrins that were in conserved co-expressed integrin-integrin pairs (e.g., ITGAV, ITGAM, and ITGB1) tended to be less disrupted by cancer than integrins (e.g., ITGA2 and ITGA3) whose integrin-integrin co-expression relationships were disrupted by cancer.

### Integrins expression patterns in metastatic breast cancer

Metastasis is the major cause of tumor-related deaths,^49^ but the biological profiles associated with metastasis, especially at distant sites, are not well understood.^50^ Recently, large scale projects, such as AURORA, have begun to investigate gene expression in metastatic samples, and we used data from AURORA to compare integrin expression in primary breast tumor with that in all metastasis sites (Fig. 6a). The expression of some integrins (e.g., ITGAX, ITGBL1, and ITGAL) was significantly lower in metastatic samples. We also created a t-SNE plot to visualize if integrin expression could distinguish primary and metastatic samples and found that there is some clustering of the metastatic samples (Fig. 6b). A Random Forest model trained to classify samples as a primary or metastatic tumors based on integrin expression had an accuracy of 0.78 (Fig. 6c). However, the F1 score for primary tumor samples was relatively low (0.65 for primary tumor and 0.84 for metastasis samples), and the model tended to improperly classify primary tumor samples as metastatic. This result may be due to sample imbalance, as there were 79 metastasis samples and only 44 primary tumor samples. The integrin-based RF model performed similarly to models based on expression of the PAM50 (mean accuracy ∼0.80, Supplementary Fig. 13a) and Donato gene groups (mean accuracy ∼0.79, Supplementary Fig. 13b).

**Fig. 6:**
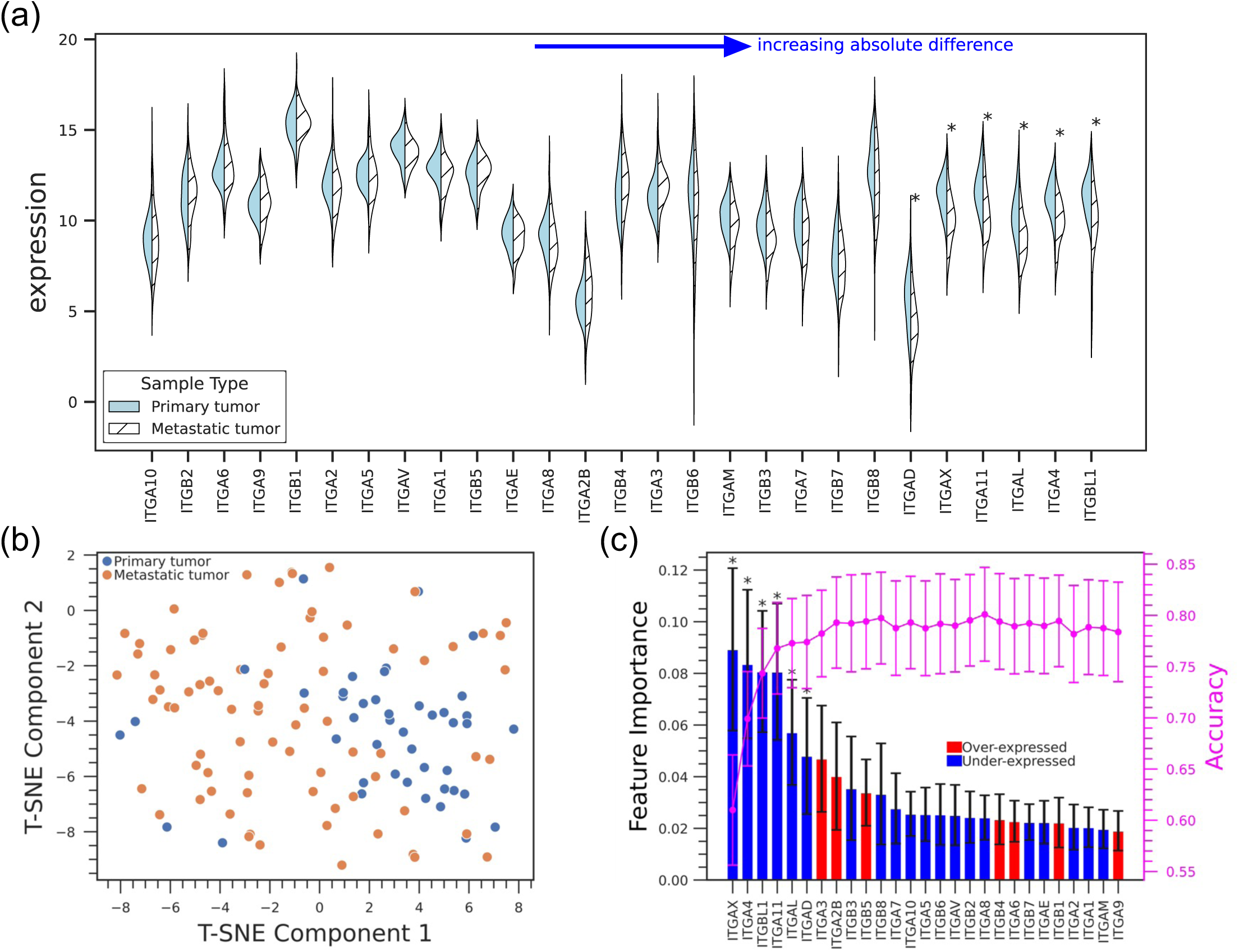
Integrin expression in metastatic breast cancer. (a) Violin plots of integrin expression in primary tumor (breast) and all metastasis sites combined from the AURORA dataset. (b) t-SNE plot based on integrin expression in AURORA (c) Feature importance plot and accuracy with increase of number of features of a Random Forest model classifying primary breast tumor and metastasis samples (accuracy is 0.78 ± 0.046, AUROC = 0.84 ± 0.041). The magenta line shows model accuracy as each feature is added from highest to lowest importance.

While the results discussed to this point compared primary tumor samples with all metastatic samples combined, we also compared integrin expression in primary tumor with samples from specific metastasis sites with the largest number of samples: liver, brain, lymph node, and lung (Fig. 7a). The difference in expression between primary samples and at least one of these metastasis sites was significant for 10 integrins, including ITGA4, ITGA11, and ITGA7 (Supplementary Table 4). In general, the expression of integrins was lower in the metastatic samples from all four of the sites (Fig. 7a), a result in agreement with expression differences between primary tumor samples and all metastatic samples (Fig. 6a).

**Fig. 7:**
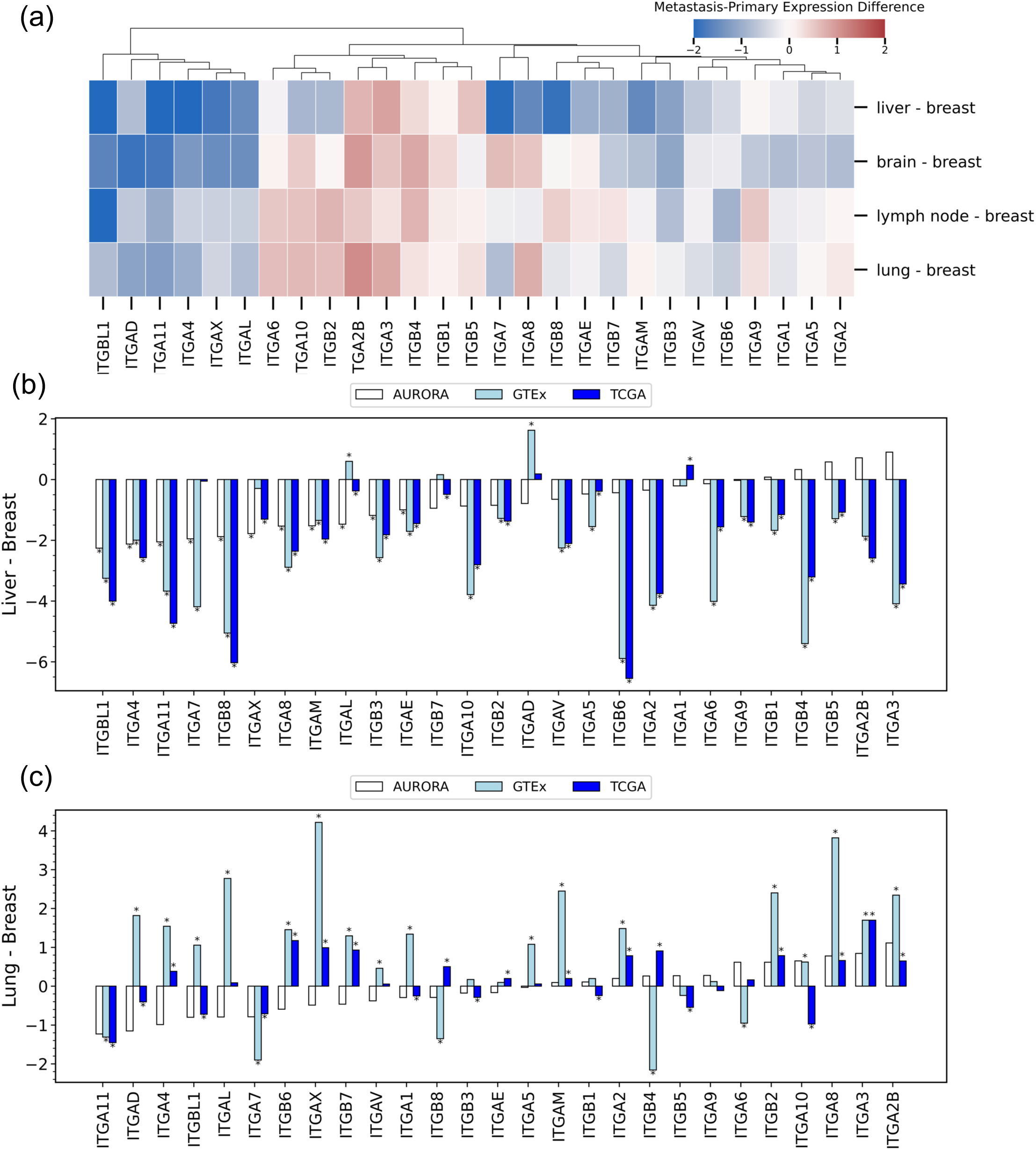
Comparison of integrin expression in breast cancer that has metastasized to different sites. (a) Heatmap of differences between the mean expression of integrins in metastatic samples (Liver, Brain, Lymph node and Lung) and in primary tumor. (b) Difference in mean expression of integrins in (metastatic) liver and primary breast tumors (AURORA), liver and breast tissue in GTEx (healthy) samples, and liver and breast (primary tumor) in TCGA samples. (c) Difference in mean expression of integrins in (metastatic) lung and primary breast tumors (AURORA), lung and breast tissue in GTEx (healthy) samples, and lung and breast (primary tumor) in TCGA samples. In (b) and (c), asterisks (*) indicate significance from independent t-test after Bonferroni correction.

The lower expression of integrins in metastasis was particularly pronounced in liver metastasis, as over 20 of the integrins had lower expression in samples from liver metastasis than in primary tumor. To put this result in context, we also compared integrin expression in breast and liver samples from the GTEx and TCGA data sets (Fig. 7b). In these comparisons, expression in liver samples was also lower than in breast samples for the majority of integrins, indicating that the difference in integrin expression in breast cancer that has metastasized to liver mirrors the differences between breast and liver in healthy and primary tumor samples. We performed a similar comparison of lung and breast tissue (Fig. 7c) and found less consistent results, as there was a mix of cases where integrin expression was higher and lower in lung tissue than breast tissue. We note that the data in Fig. 7b and c can be used to compare changes in integrin expression (i.e. whether expression was higher or lower in liver/lung than in breast) but should not be used to compare the magnitudes of these changes as the AURORA and GTEx/TCGA data were not normalized together.

## Discussion

Integrins have long been established as key players in cancer, but the specifics of relationships between integrin expression patterns and cancer and how they vary across tissues and cancer types have yet to be fully elucidated. In this work, we attempted to unravel some of these relationships by using machine learning methods to investigate integrin expression patterns in healthy and cancer samples from 8 solid tissues. We found that integrin expression patterns had sufficient variation across tissues and cancer types to enable the creation of Random Forest models that could classify samples by tissue/cancer type or as healthy/cancer using integrin expression alone. A wide array of integrins were important features in these classifiers, and, for most tissues, different integrins were the important features for healthy tissues and their corresponding cancers. There were a couple of exceptions to this trend, as ITGA8 and ITGA2B were the most important integrins in the classification of both healthy and cancer samples in prostate and testis, respectively, and ITGB4 was important in the classification of both healthy and cancer samples in stomach and colon.

We focused on breast cancer and investigated the impact of cancer development on the co-expression relationships of integrins and how integrin expression patterns change during metastasis. The co-expression of most integrins was significantly altered in breast tissue after cancer development, but there was a group of integrins, including ITGAM and ITGB2, whose co-expression relationships were largely conserved between healthy and cancer breast samples. Metastasis was shown to result in significantly lower expression of some integrins, including ITGA4, ITGA11, and ITGA7. This reduced expression in metastasis was particularly notable for metastasis to the liver, behavior similar to the lower expression of integrins in liver and liver cancer in comparison with healthy breast and breast cancer, respectively.

One of the major results of this work is that, with some exceptions (e.g., the reduced expression of integrins in pancreatic cancer compared to healthy pancreas tissue), the expression of integrins cannot be considered as a unified block. Instead, the behavior of individual integrins must be considered. To this end, we summarized important results identified in this work for each integrin (Supplemental Table 5). Some examples of specific integrins whose expression had interesting behavior include:

### ITGA7

The RF models in this work identified ITGA7 as the key integrin that enabled the classification of breast tissue samples as healthy or cancer, corresponding with previous studies that identified ITGA7 as a potential predictive marker of chemotherapy response and found that that it was involved in regulating migration and invasion in breast cancer.^19,43^ The expression of ITGA7 was found to be lower in breast cancer in comparison with corresponding healthy tissues, a result that may be explained by previous findings that the ITGA7 gene promoter CpG island, along with promoter regions of several other integrins (e.g., ITGA1, ITGA4), are abnormally hypermethylated in breast cancer samples.^51^ Reduced expression of ITGA7 in cancer was not unique to breast, as its expression was lower in most cancers in comparison with corresponding healthy tissues. ITGA7 was also found to be the most important integrin when classifying samples as healthy or cancer in liver, lung, and prostate (Supplementary Fig. 9). Furthermore, ITGA7 had significantly lower expression in breast cancer that metastasized to liver than in primary breast tumors.

### ITGA3

Associations between expression of ITGA3 and cancer risk are complex, as high ITGA3 expression has been associated with an increased risk in some cancers and a reduced risk in others.^14,15^ In breast cancer, for example, ITGA3 was shown to have a methylated promoter region, suggesting gene silencing, and high ITGA3 expression was found to be associated with improved relapse-free survival (RFS) among breast cancer patients.^15^ Interestingly, ITGA3 was identified as the integrin with the highest feature importance in both the multi-class RF model that classified cancer samples by cancer type and the one-vs-all classifiers for breast and lung (LUAD) cancers (Fig. 2 and Table 1). The importance of ITGA3 in classifying samples was unique to cancers, as it was not highly ranked in any models that classified healthy samples by tissue. The difference in the importance of ITGA3 in classifying healthy and cancer samples was relatively surprising given the relatively modest differences between mean ITGA3 expression in healthy tissues and their corresponding cancers (Fig. 3). However, we did find that cancer development had a strong impact on the co-expression relationships of ITGA3 in breast samples (Fig. 5 and Supplementary Tables 2 and 3).

### ITGAV

ITGAV has been investigated as a therapeutic target in cancers for over 20 years.^52,53^ However, these studies have yet to translate to therapeutic benefits for patients, and it has been suggested that future targeting of ITGAV containing heterodimers, such as *αvβ3*, relies on understanding their expression in individual tumors.^53^ For the tissues/cancers investigated in this work, the expression of ITGAV was relatively consistent across tissues and healthy/cancer conditions, and ITGAV was not identified as an integrin with a high feature importance in any of the RF models that classified samples by tissue or disease status. Additionally, its co-expression was relatively well conserved across healthy and cancer samples when considering both its co-expression with other integrins (i.e., 2 of the 4 conserved integrin-integrin co-expression relationships involved ITGAV) and with all protein coding genes (Supplementary Fig. 11).

### ITGB4

ITGB4 has been reported to be up-regulated in colon cancer and associated with overall survival, giving it a potential role as both a therapeutic target and prognostic marker for this cancer type.^54^ The analysis in this work also showed that the expression of ITGB4 increased in colon cancer and, furthermore, identified ITGB4 as the second most important feature in distinguishing both healthy colon and colon cancer samples from other healthy tissues/cancer. Additionally, ITGB4 was the integrin that had the largest increase in expression in lung cancer compared to healthy lung tissue, corresponding with previous studies that identified ITGB4 as a potential candidate marker for tumor status and a prognostic indicator of small cell lung carcinoma (SCLC).^55^

While this study was able to characterize integrin expression patterns across many tissues and their corresponding cancers, we recognize that it suffers from several limitations and presents opportunities for future analysis. First, our analysis focused on bulk RNA-Seq datasets and did not consider single-cell expression. Therefore, we cannot determine the extent to which the observed differences in expression patterns are due to differences in the mixtures of cell types in different samples. Second, this study analyzed only mRNA expression and did not include measures of integrin protein amounts in cells or in functional heterodimers on cell surfaces. However, we believe that the results presented here can be used to create hypotheses for future bioinformatic and experimental analysis. As the number of single-cell and expression datasets continues to grow, we plan on using these data sets to continue to investigate relationships between integrin expression patterns and cancer.

## Acknowledgements

The authors wish to thank Dr Janusz Rak at McGill University for inspiring this study and providing insightful comments.

## Author Contributions

The authors wish it be known that H.S. and S.G. contributed equally and are joint first authors. Co-first authors can prioritize their own names when citing or referring to this paper in their resume/portfolio. J.D.Z., X.H., and Y.W. initially conceptualized the study. H.S. and S.G. conducted all analysis, wrote python code and prepared all figures and tables, with input from Y.W., Y.J., Q.C., and J.D.Z. H.S. and S.G. conducted all statistical analysis with input from Y.W., Y.J., and J.D.Z. H.S. and S.G. prepared first draft of manuscript and H.S., S.G., X.H., Y.J., Q.C., J.D.Z. and Y.W. edited the draft.

## Data Availability

This study is based on publicly available datasets (GTEx, TCGA, AURORA-US).

