## Supplemental Files for "A Machine Learning-Based Investigation of Integrin Expression Patterns in Cancer and Metastasis"

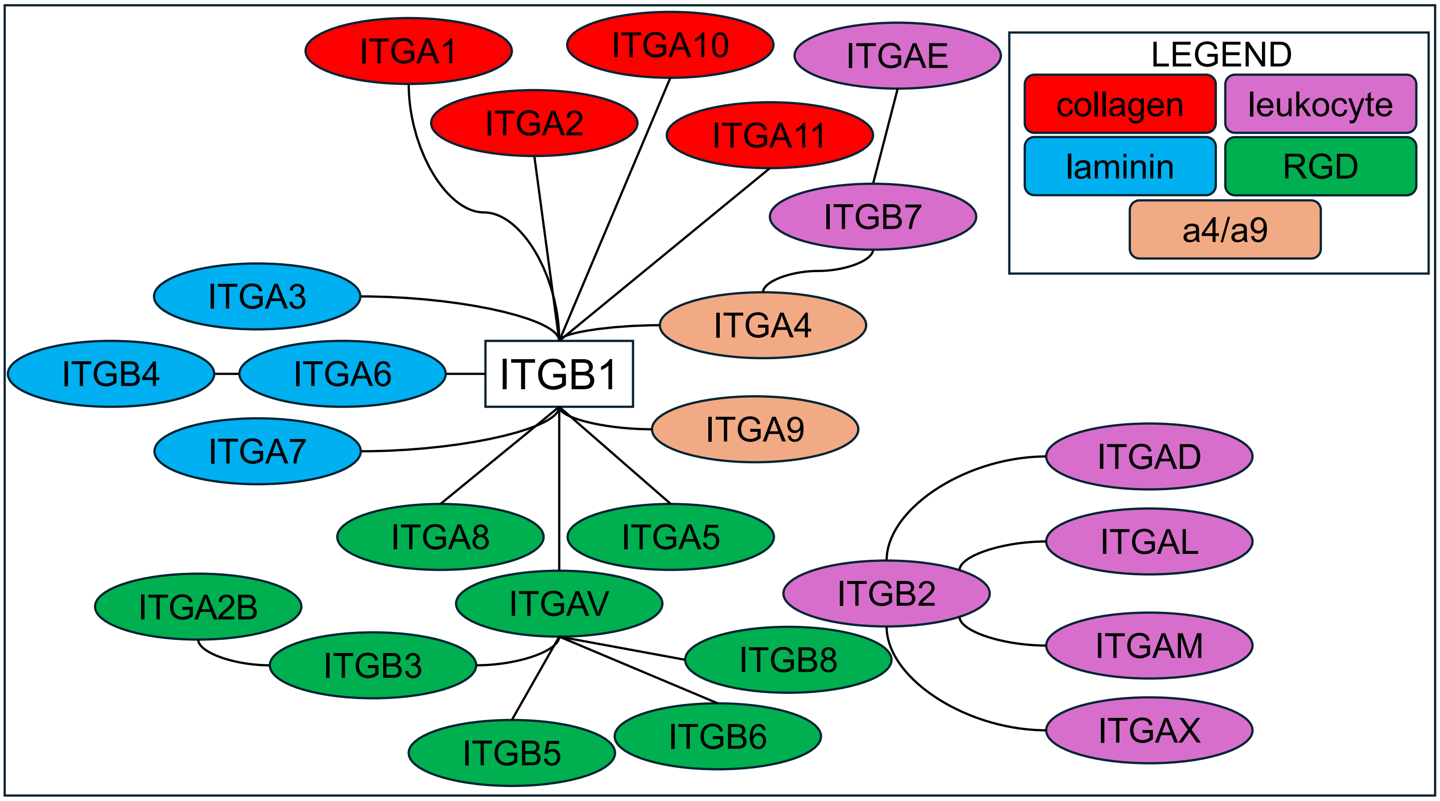


**Supplementary Figure 1:** Known integrin heterodimers and subunit classifications. ITGB1, shown in white, dimerizes with α-subunits from most classes, while the other integrins are colored by class as shown in the figure legend.

**Supplementary Table 1:** Sample sizes of GTEx tissues and corresponding TCGA cancer types[*]

| GTEx tissue | Samples | TCGA cancer type | Primary tumor samples | Patient matched normal/tumor samples |
|---|---|---|---|---|
| Breast[**] | 179 | Breast Invasive Carcinoma (BRCA) | 1092 | 112 |
| Prostate | 100 | Prostate Adenocarcinoma (PRAD) | 495 | 52 |
| Liver | 110 | Liver Hepatocellular Carcinoma (LIHC) | 369 | 50 |
| Lung | 288 | Lung Adenocarcinoma (LUAD) | 513 | 58 |
| | | Lung Squamous Cell Carcinoma (LUSC) | 498 | 50 |
| Pancreas | 167 | Pancreatic Adenocarcinoma (PAAD) | 178 | 4 |
| Stomach | 174 | Stomach Adenocarcinoma (STAD) | 414 | 33 |
| Colon – Transverse | 167 | Colon Adenocarcinoma (COAD) | 288 | 26 |
| Colon – Sigmoid | 141 | | | |
| Testis | 165 | Testicular Germ Cell Tumor (TGCT) | 148 | 0 |

[*]For GTEx colon and TCGA lung data, two different tissue/cancer types were included. Unless otherwise specified, GTEx samples for sigmoid and transverse regions of the colon were combined and treated as healthy colon tissue, while lung adenocarcinoma and lung squamous carcinoma samples TCGA were treated as distinct cancer types. There were patient-matched normal tissue samples for some cancer types used in some analysis in the study, as described in the appropriate sections.

[**]For some analysis of breast tissue (Random Forest models that classified samples as healthy or cancer breast tissue and co-expression analysis), we selected only samples from female donors, resulting in 80 GTEx healthy samples, 1092 BRCA primary tumor samples, and 111 tumor/normal BRCA patient matched samples.

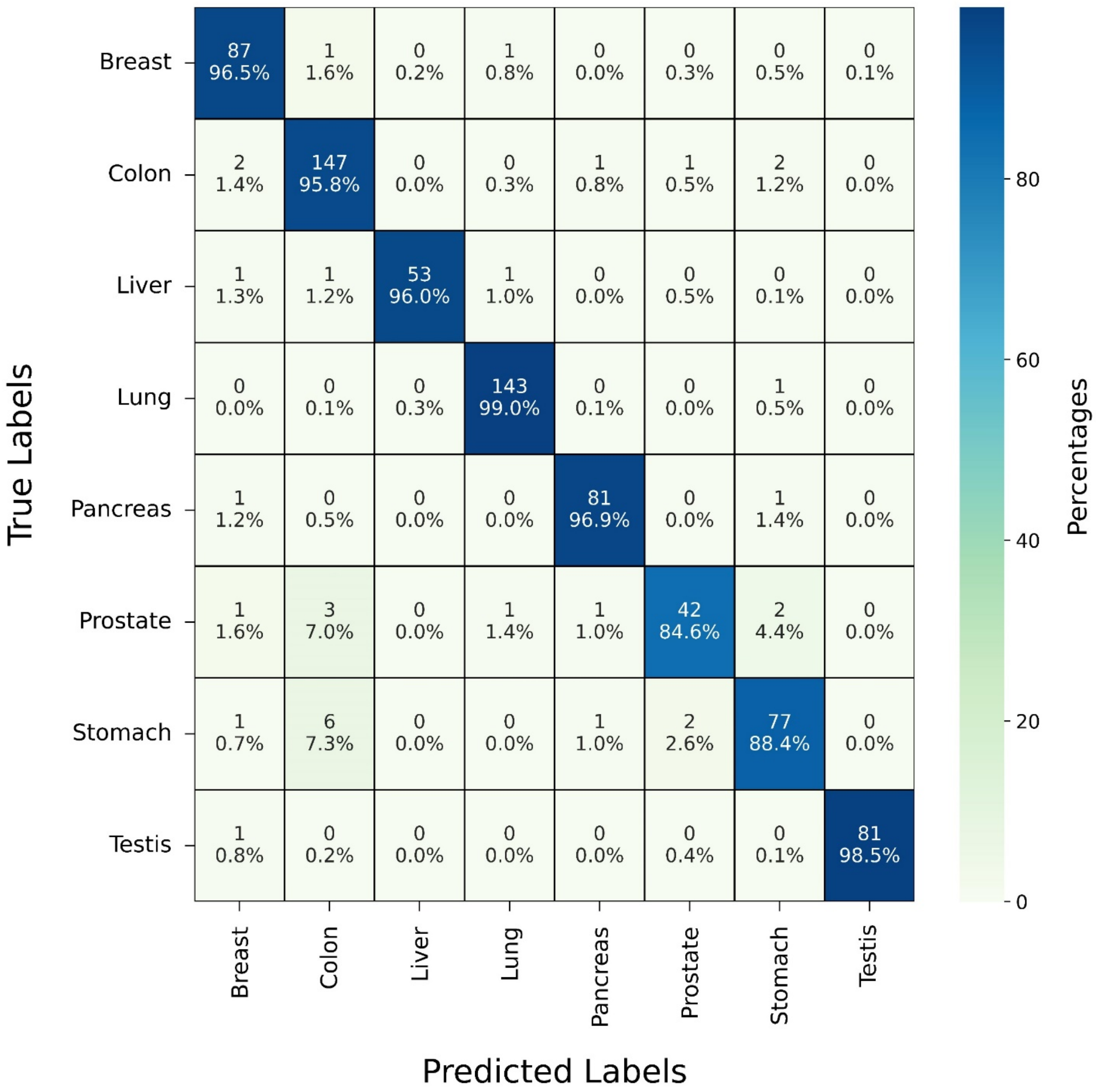


**Supplementary Figure 2:** Confusion matrix of multiclass Random Forest model that classifies healthy (GTEx) samples by tissue type based on integrin expression. Each cell shows the average over 500 Random Forest trials of the counts of predictions and the percentage of predicted labels normalized by the number of true labels for each row. The color scale is based on the percentage of true predictions.

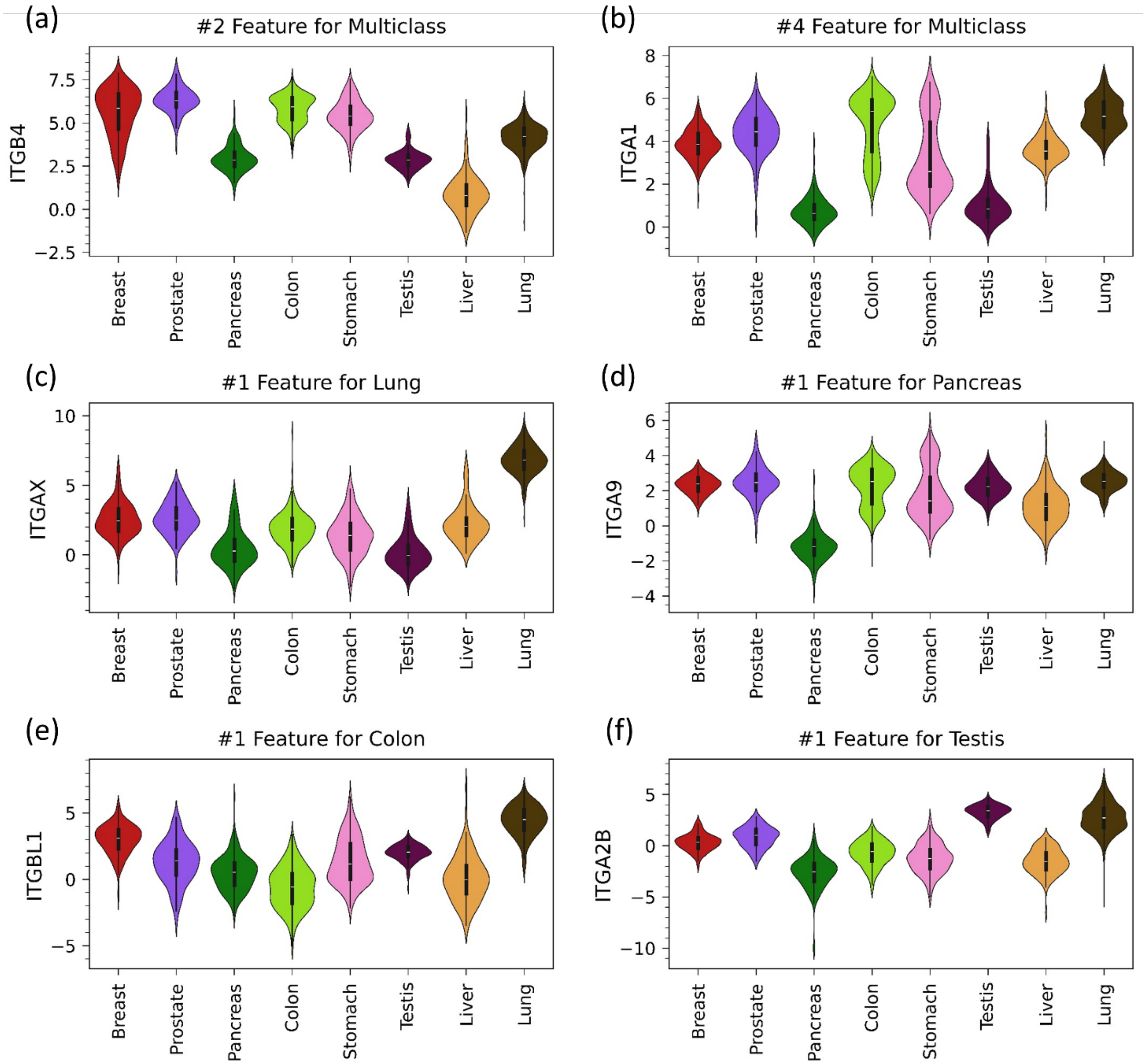


**Supplementary Figure 3:** Violin plots of mRNA expression in healthy tissues of the integrins with the highest feature importance in multiclass and one-vs-all Random Forest classifiers of samples by healthy tissue type. The classifier in which the integrin had high importance and its rank among features in that model are given in the title of each panel.

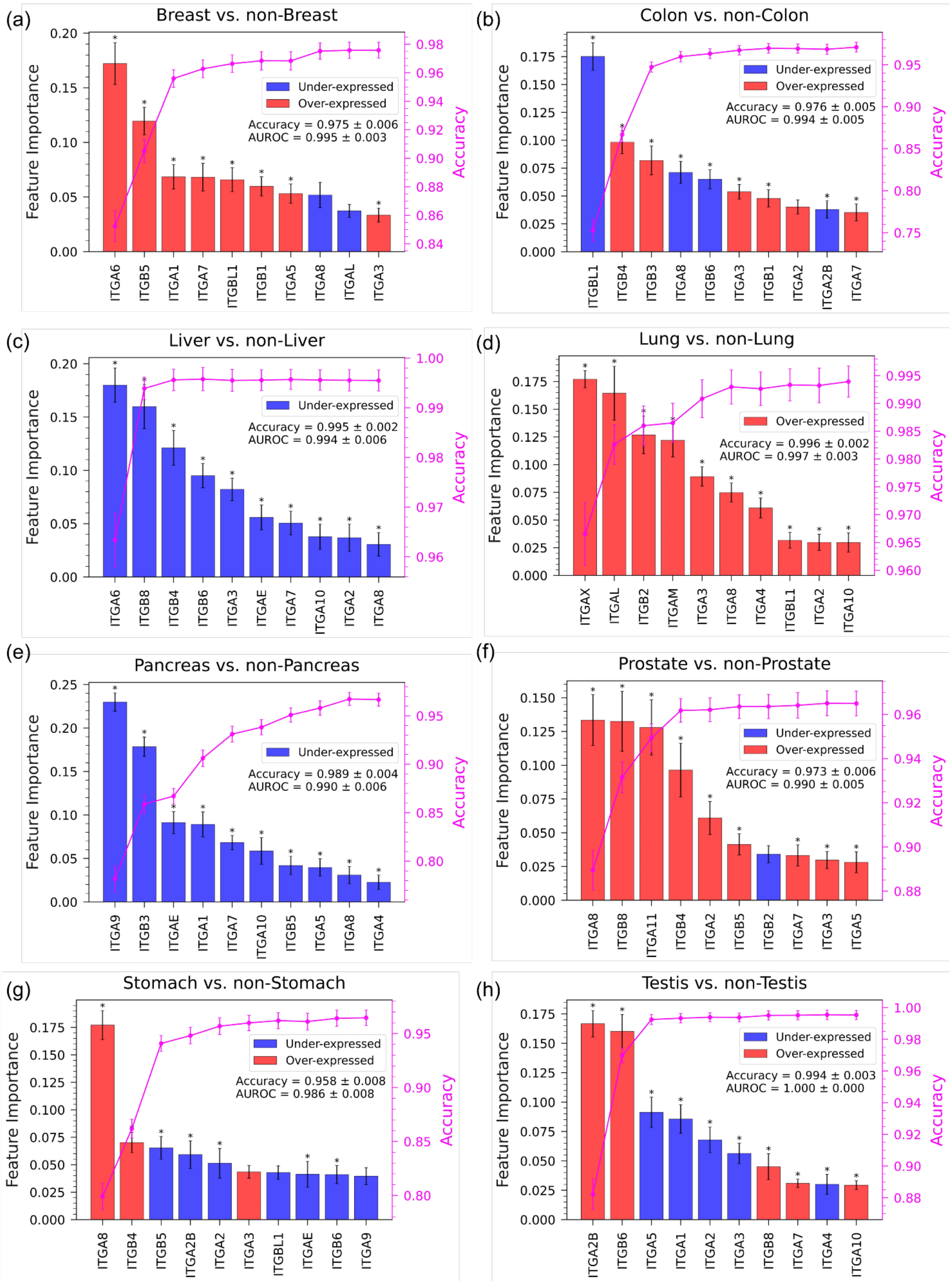


**Supplementary Figure 4.** One-vs-all Random Forest classifiers of healthy (GTEx) tissue samples classified by expression. Each plot shows the feature importance of the top ten integrin sin a model trained using all integrins and the increase in accuracy with the number of integrins added to the model averaged over 500 trials. The examined tissue type for each panel is indicated in the panel title. The reported accuracy and AUROC on each panel are averages over 500 trials of the model using 27 integrins.

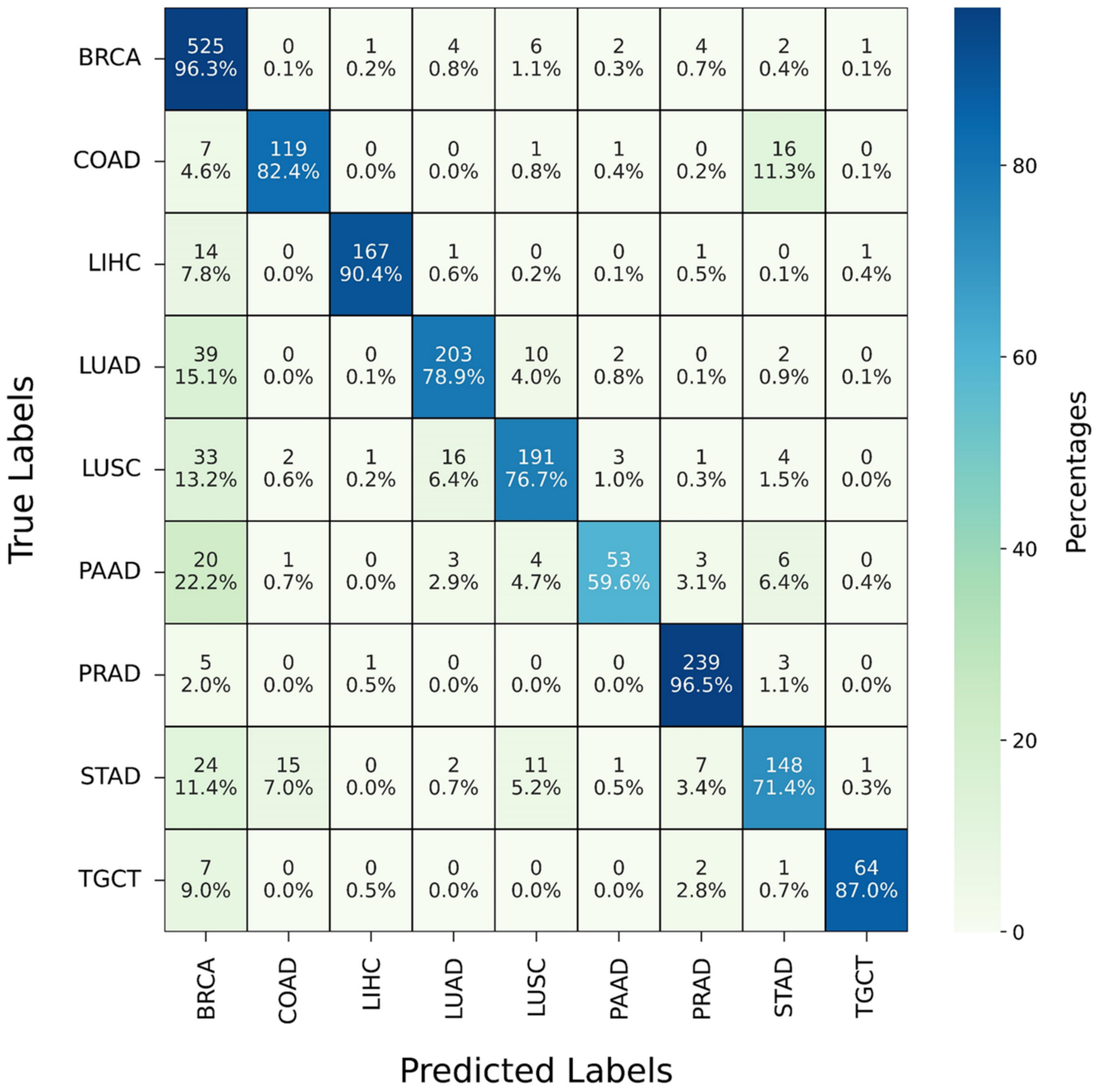


**Supplementary Figure 5:** Confusion matrix of a multiclass Random Forest model that classifies cancer (TCGA) samples by tissue type based on the expression of 27 integrins. Each cell shows the average over 500 Random Forest trials of the number of counts of predictions and the percentage of predicted labels normalized by the number of true labels for each row. The color scale is based on the percentage of true predictions.

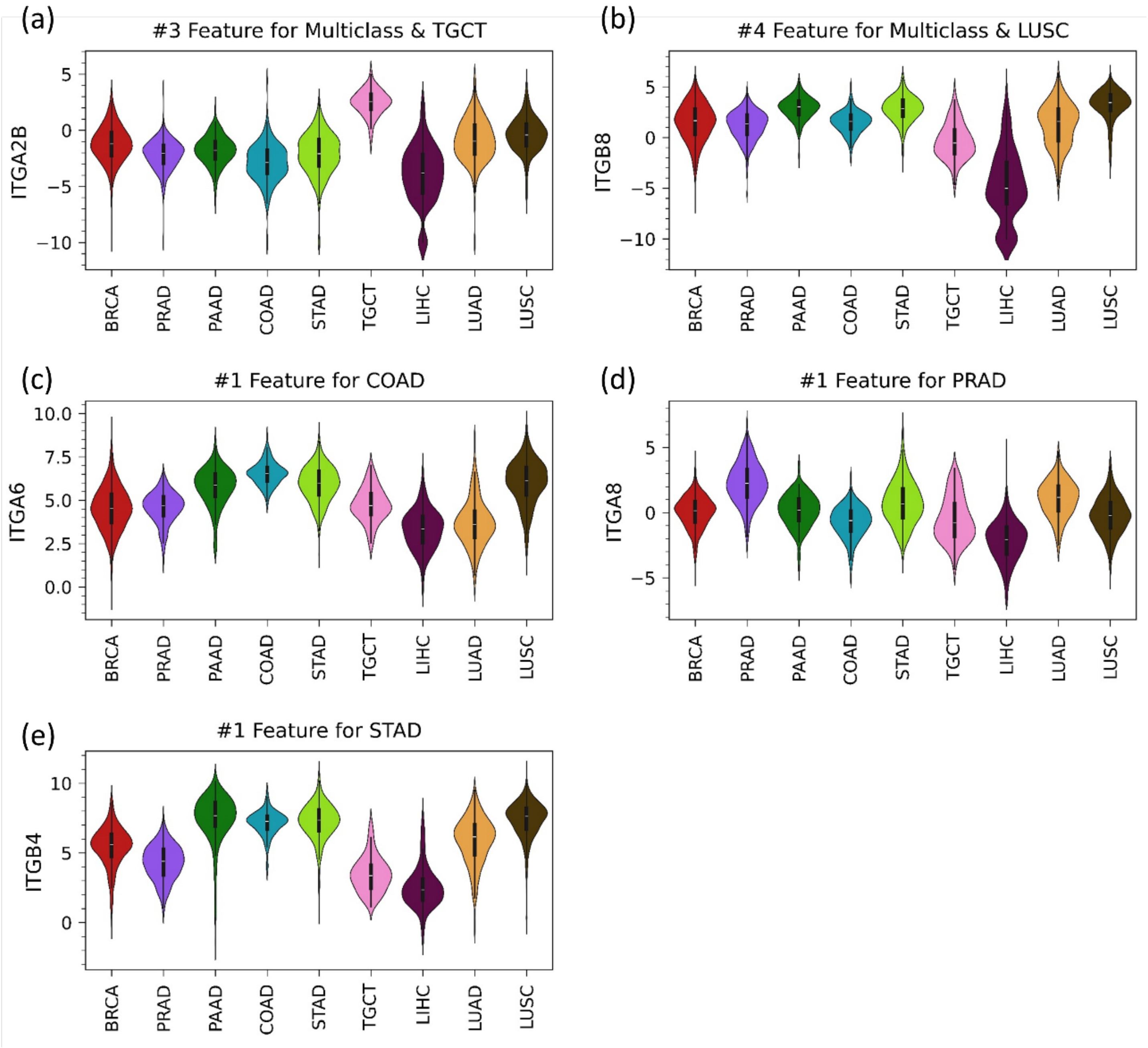


**Supplementary Figure 6:** Violin plots of mRNA expression in TCGA primary tumors of the integrins with the highest feature importance in multiclass and one-vs-all Random Forest classifiers of samples by cancer type. The classifier in which the integrin had high importance and its rank among features in that model are given in the title of each panel.

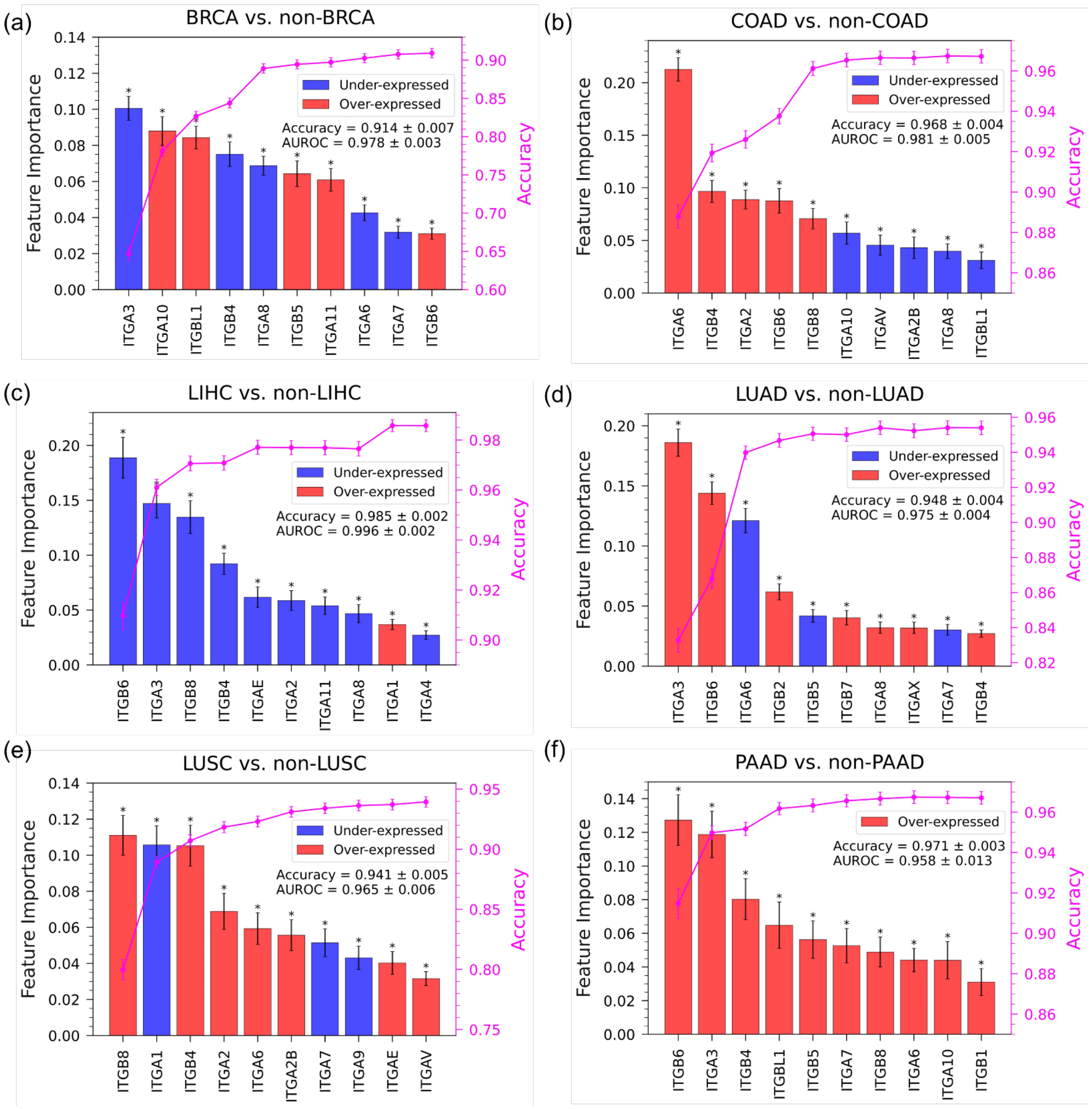

(a)
BRCA vs. non-BRCA
Under-expressed
Over-expressed
Accuracy = 0.914 ± 0.007
AUROC = 0.978 ± 0.003
Feature Importance
Accuracy
ITGA3 ITGA10 ITGBL1 ITGB4 ITGA8 ITGB5 ITGA11 ITGA6 ITGA7 ITGB6
(b)
COAD vs. non-COAD
Under-expressed
Over-expressed
Accuracy = 0.968 ± 0.004
AUROC = 0.981 ± 0.005
Feature Importance
Accuracy
ITGA6 ITGB4 ITGA2 ITGB6 ITGB8 ITGA10 ITGAV ITGA2B ITGA8 ITGBL1
(c)
LIHC vs. non-LIHC
Under-expressed
Over-expressed
Accuracy = 0.985 ± 0.002
AUROC = 0.996 ± 0.002
Feature Importance
Accuracy
ITGB6 ITGA3 ITGB8 ITGB4 ITGAE ITGA2 ITGA11 ITGA8 ITGA1 ITGA4
(d)
LUAD vs. non-LUAD
Under-expressed
Over-expressed
Accuracy = 0.948 ± 0.004
AUROC = 0.975 ± 0.004
Feature Importance
Accuracy
ITGA3 ITGB6 ITGA6 ITGB2 ITGB5 ITGB7 ITGA8 ITGAX ITGA7 ITGB4
(e)
LUSC vs. non-LUSC
Under-expressed
Over-expressed
Accuracy = 0.941 ± 0.005
AUROC = 0.965 ± 0.006
Feature Importance
Accuracy
ITGB8 ITGA1 ITGB4 ITGA2 ITGA6 ITGA2B ITGA7 ITGA9 ITGAE ITGAV
(f)
PAAD vs. non-PAAD
Over-expressed
Accuracy = 0.971 ± 0.003
AUROC = 0.958 ± 0.013
Feature Importance
Accuracy
ITGB6 ITGA3 ITGB4 ITGBL1 ITGB5 ITGA7 ITGB8 ITGA6 ITGA10 ITGB1

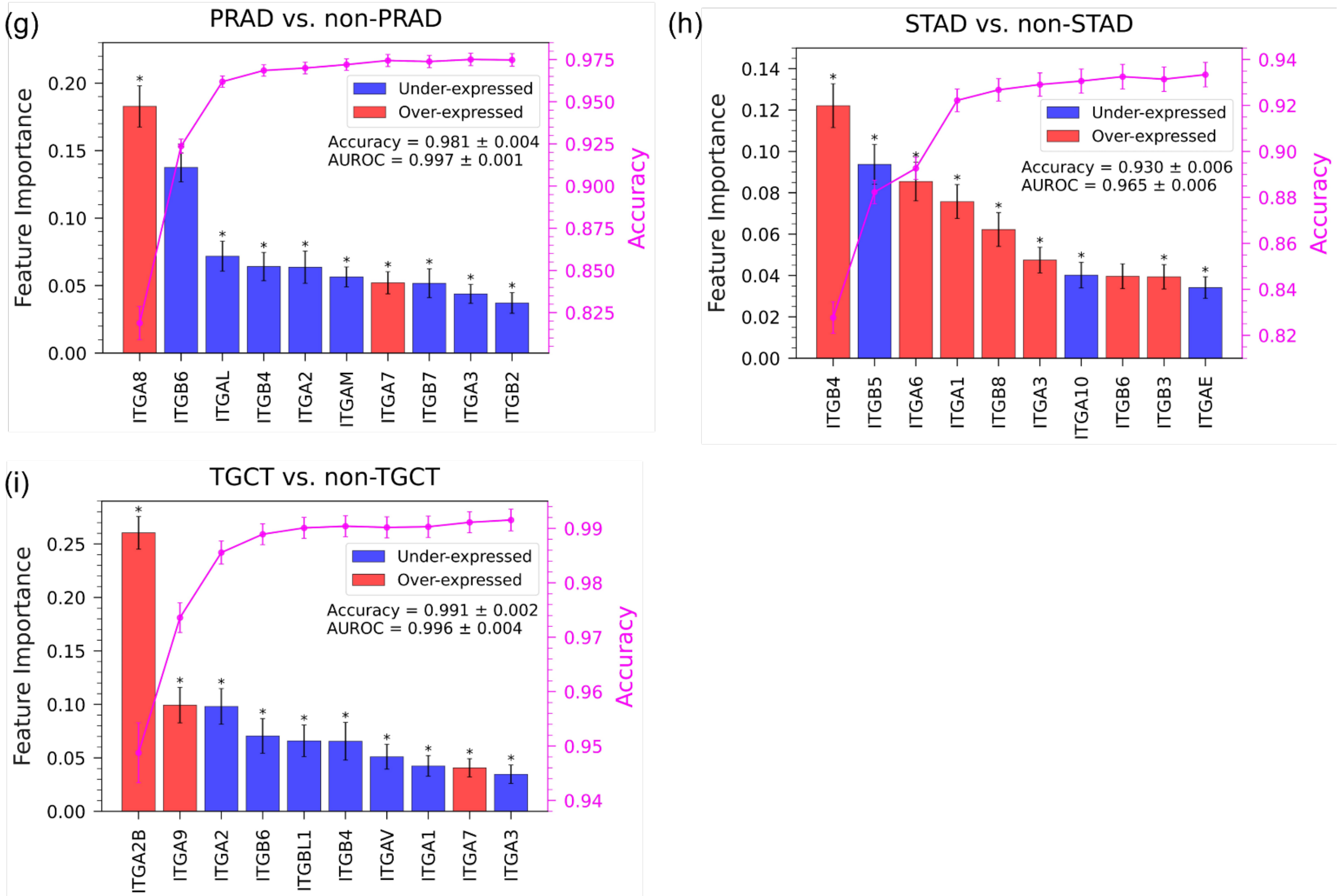


**Supplementary Figure 7:** One-vs-all Random Forest classifiers of cancer (TCGA) tissue samples classified by integrin expression. Each plot shows the feature importance of the top ten integrins in a model trained using all integrins and the increase in accuracy with the number of integrins added to the model averaged over 500 trials. The examined tissue type for each panel is indicated in the panel title. The reported accuracy and AUROC on each panel are averages over 500 trials of the model using 27 integrins.

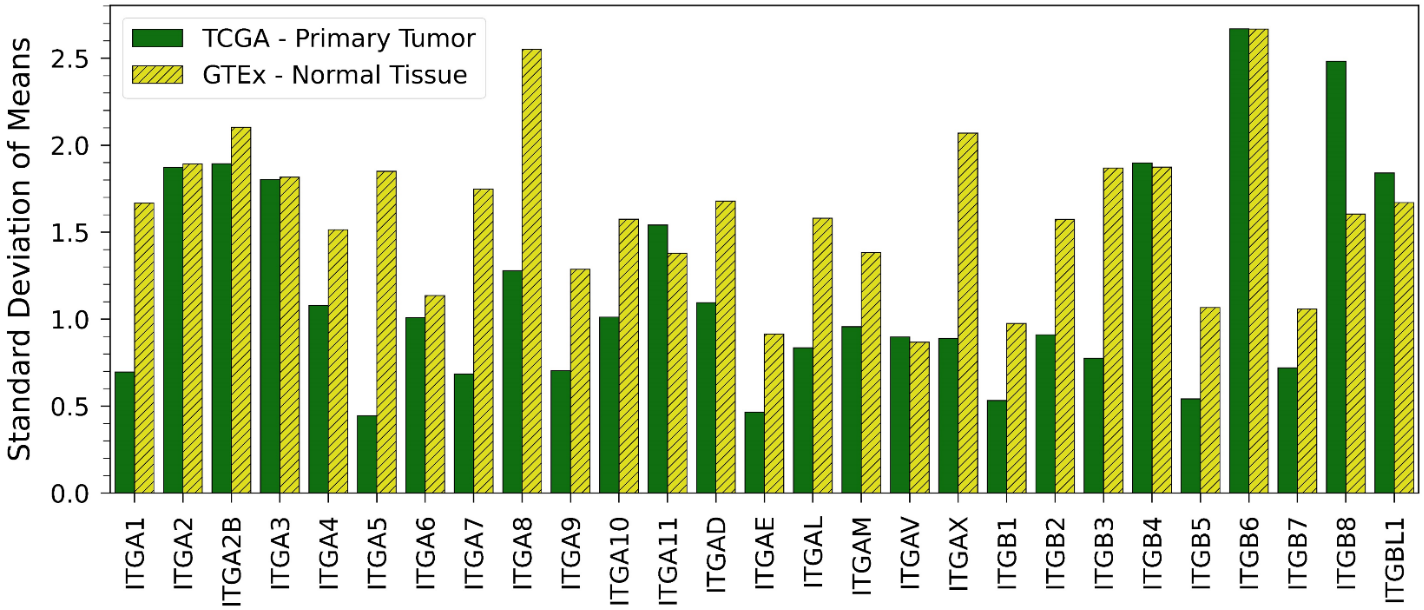


**Supplementary Figure 8:** Standard deviation in integrin expression for GTEx healthy tissues and TCGA primary tumors. The standard deviations were calculated from the mean expression in each tissue/cancer type given in Supplementary Table 1.

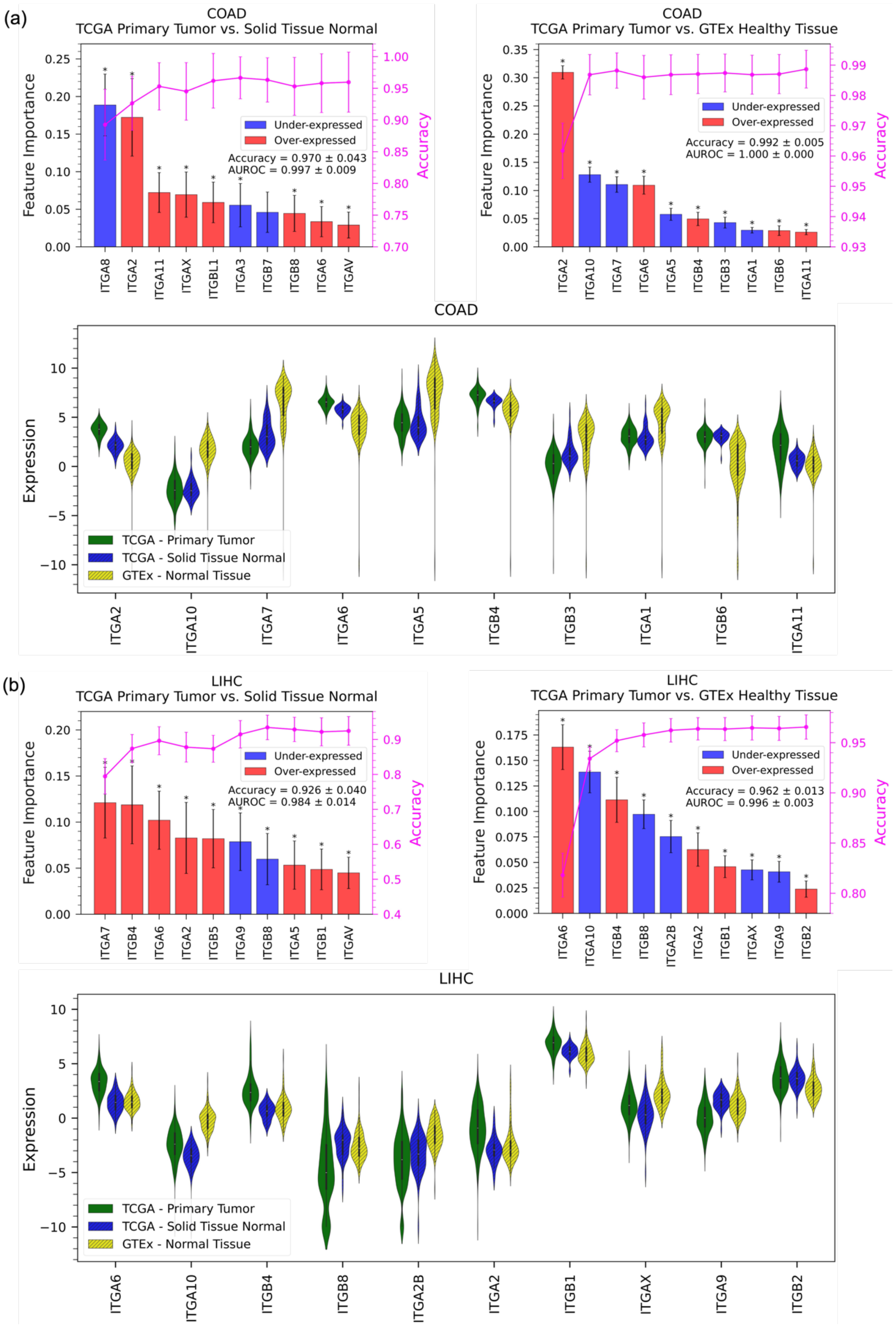
(a)
COAD
TCGA Primary Tumor vs. Solid Tissue Normal
Feature Importance
Accuracy
Under-expressed
Over-expressed
Accuracy = 0.970 ± 0.043
AUROC = 0.997 ± 0.009
ITGA8
ITGA2
ITGA11
ITGAX
ITGBL1
ITGA3
ITGB7
ITGB8
ITGA6
ITGAV
COAD
TCGA Primary Tumor vs. GTEx Healthy Tissue
Feature Importance
Accuracy
Under-expressed
Over-expressed
Accuracy = 0.992 ± 0.005
AUROC = 1.000 ± 0.000
ITGA2
ITGA10
ITGA7
ITGA6
ITGA5
ITGB4
ITGB3
ITGA1
ITGB6
ITGA11
COAD
Expression
TCGA - Primary Tumor
TCGA - Solid Tissue Normal
GTEx - Normal Tissue
ITGA2
ITGA10
ITGA7
ITGA6
ITGA5
ITGB4
ITGB3
ITGA1
ITGB6
ITGA11
(b)
LIHC
TCGA Primary Tumor vs. Solid Tissue Normal
Feature Importance
Accuracy
Under-expressed
Over-expressed
Accuracy = 0.926 ± 0.040
AUROC = 0.984 ± 0.014
ITGA7
ITGB4
ITGA6
ITGA2
ITGB5
ITGA9
ITGB8
ITGA5
ITGB1
ITGAV
LIHC
TCGA Primary Tumor vs. GTEx Healthy Tissue
Feature Importance
Accuracy
Under-expressed
Over-expressed
Accuracy = 0.962 ± 0.013
AUROC = 0.996 ± 0.003
ITGA6
ITGA10
ITGB4
ITGB8
ITGA2B
ITGA2
ITGB1
ITGAX
ITGA9
ITGB2
LIHC
Expression
TCGA - Primary Tumor
TCGA - Solid Tissue Normal
GTEx - Normal Tissue
ITGA6
ITGA10
ITGB4
ITGB8
ITGA2B
ITGA2
ITGB1
ITGAX
ITGA9
ITGB2

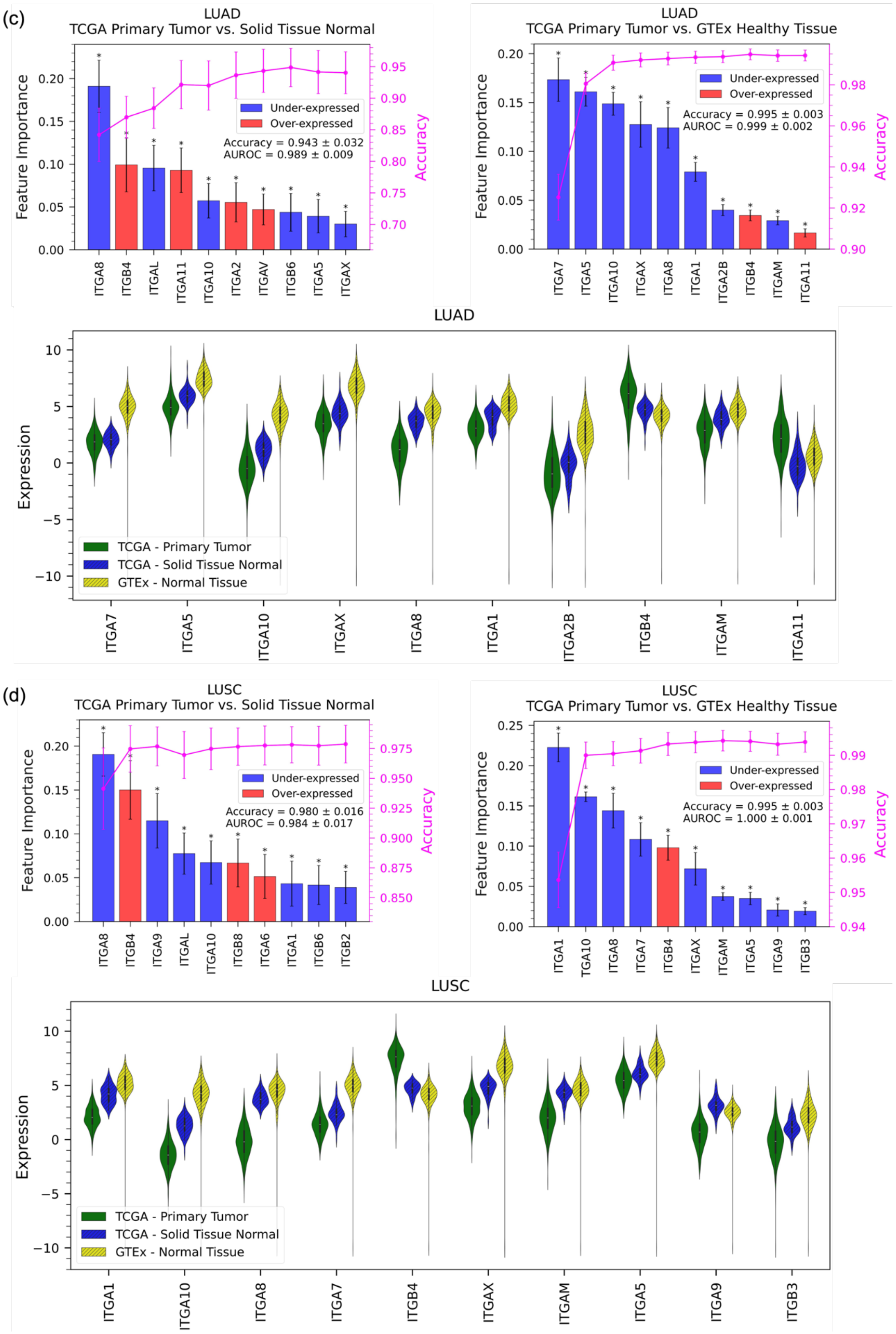

(c)
LUAD
TCGA Primary Tumor vs. Solid Tissue Normal
Feature Importance
Accuracy
Under-expressed
Over-expressed
Accuracy = 0.943 ± 0.032
AUROC = 0.989 ± 0.009
ITGA8
ITGB4
ITGAL
ITGA11
ITGA10
ITGA2
ITGAV
ITGB6
ITGA5
ITGAX
LUAD
TCGA Primary Tumor vs. GTEx Healthy Tissue
Feature Importance
Accuracy
Under-expressed
Over-expressed
Accuracy = 0.995 ± 0.003
AUROC = 0.999 ± 0.002
ITGA7
ITGA5
ITGA10
ITGAX
ITGA8
ITGA1
ITGA2B
ITGB4
ITGAM
ITGA11
LUAD
Expression
TCGA - Primary Tumor
TCGA - Solid Tissue Normal
GTEx - Normal Tissue
ITGA7
ITGA5
ITGA10
ITGAX
ITGA8
ITGA1
ITGA2B
ITGB4
ITGAM
ITGA11
(d)
LUSC
TCGA Primary Tumor vs. Solid Tissue Normal
Feature Importance
Accuracy
Under-expressed
Over-expressed
Accuracy = 0.980 ± 0.016
AUROC = 0.984 ± 0.017
ITGA8
ITGB4
ITGA9
ITGAL
ITGA10
ITGB8
ITGA6
ITGA1
ITGB6
ITGB2
LUSC
TCGA Primary Tumor vs. GTEx Healthy Tissue
Feature Importance
Accuracy
Under-expressed
Over-expressed
Accuracy = 0.995 ± 0.003
AUROC = 1.000 ± 0.001
ITGA1
TGA10
ITGA8
ITGA7
ITGB4
ITGAX
ITGAM
ITGA5
ITGA9
ITGB3
LUSC
Expression
TCGA - Primary Tumor
TCGA - Solid Tissue Normal
GTEx - Normal Tissue
ITGA1
ITGA10
ITGA8
ITGA7
ITGB4
ITGAX
ITGAM
ITGA5
ITGA9
ITGB3

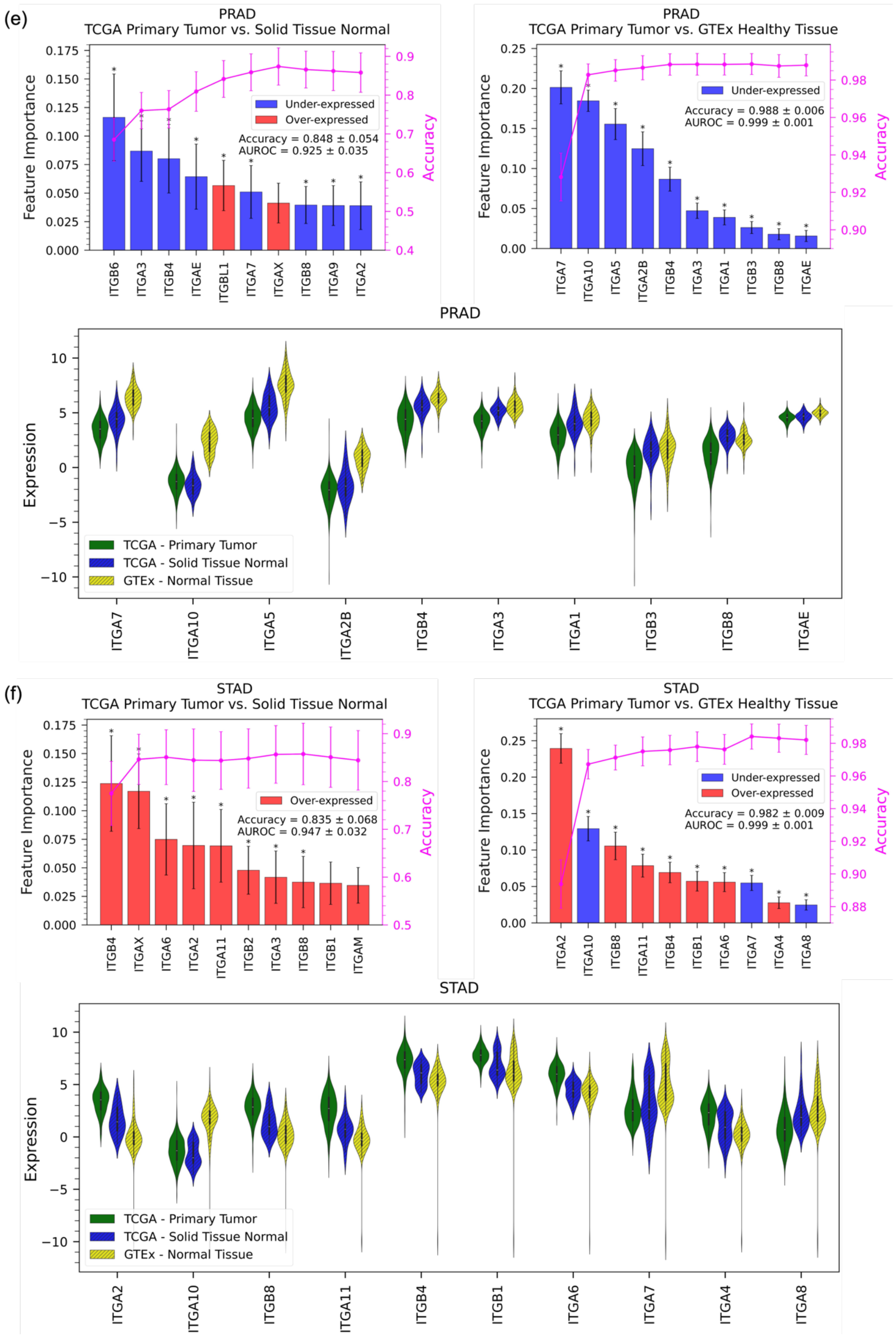


**Supplementary Figure 9:** Random Forest model results trained/tested on TCGA patient-matched normal/tumor dataset (left column) and TCGA tumor/GTEx heathly tissues for other types of tissues/tumors dicussed in this project. The reported accuracy and AUROC on each panel are averages over 500 trials of the model using 27 integrins. The violin plots show the expression of top 10 integrins identified by TCGA primary tumor vs. GTEx healthy tissues RF model.

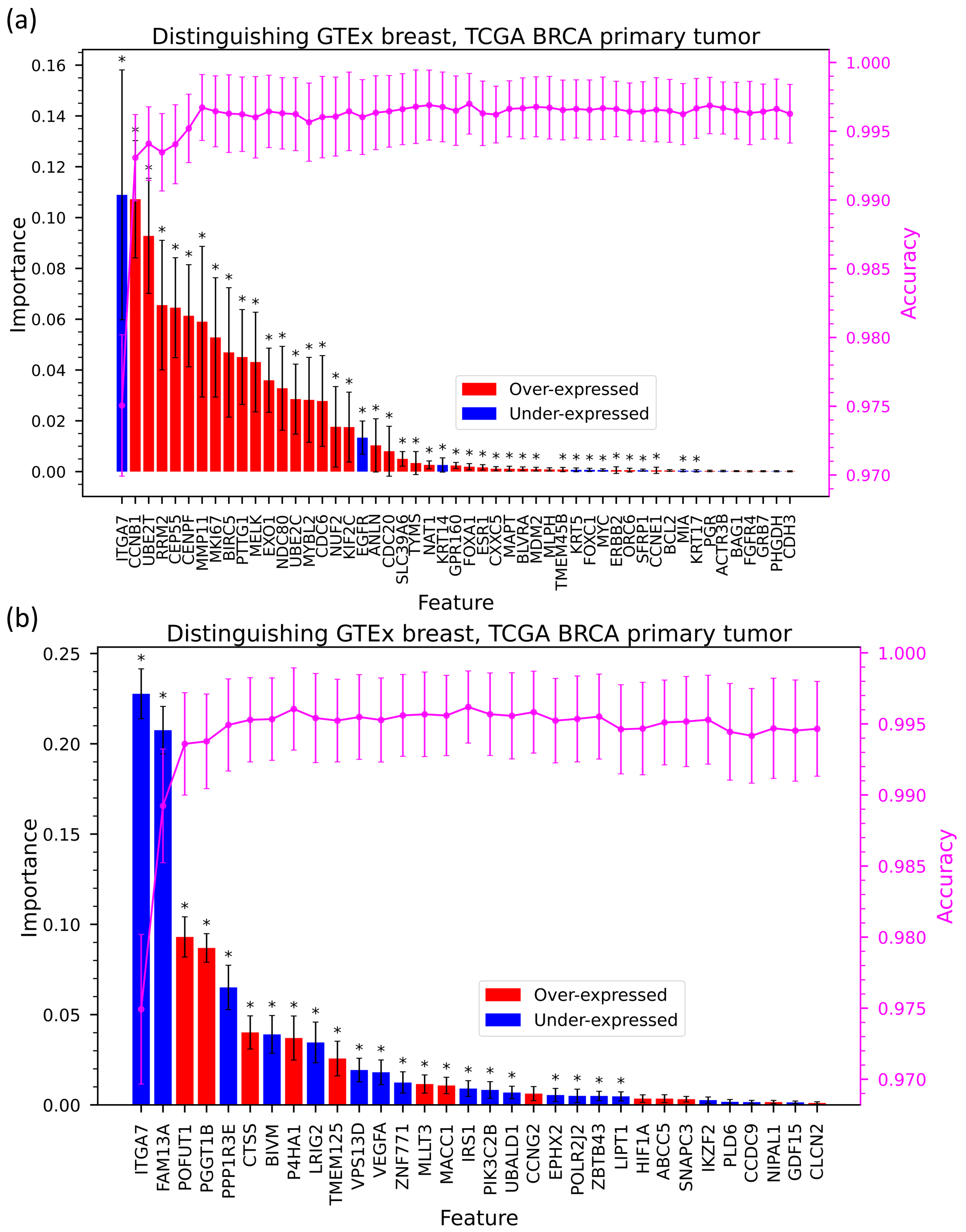


**Supplementary Fig. 10:** (a) Feature importance in a Random Forest model using a 50%/50% training/test split that classifies breast samples as healthy or cancer using the PAM50 gene set and ITGA7 as features. The magenta line shows the change in the accuracy of the model as features are added one-by-one from the most important feature to the least. The accuracy of the model using all features is 1.00 ± 0.0022 and AUROC is 1.00 ± 0.004. (b) Feature importance in a Random Forest model using a 50%/50% training/test split that classifies breast samples as healthy or cancer using a gene set identified by Donato *et al.* (CTC gene sets, 32 genes) and ITGA7 as features. The magenta line shows the change in the accuracy of the model as features are added one-by-one from the most important feature to the least. The accuracy of the model using all features is 0.99±0.0032 and AUROC is 1.000 ± 0.001.

**Supplementary Table 2:** Pearson correlation coefficients for integrin-integrin pairs that are highly correlated (Pearson R ≥ 0.6) in GTEx, and their corresponding coefficients in TCGA BRCA primary tumor samples*

| Integrin | Integrin | GTEx R | TCGA Primary Tumor R | STRING-db** Association? |
|---|---|---|---|---|
| ITGA2 | ITGB8 | 0.87 | 0.34 | A, C, D |
| ITGB4 | ITGB8 | 0.87 | 0.34 | None |
| ITGA2 | ITGA3 | 0.84 | 0.41 | A, C, D |
| ITGA2 | ITGB6 | 0.83 | 0.41 | A, C, D |
| ITGB6 | ITGB8 | 0.81 | 0.25 | None |
| ITGA2 | ITGB4 | 0.79 | 0.12 | None |
| ITGAL | ITGA4 | 0.76 | 0.56 | None |
| ITGAL | ITGB7 | 0.76 | 0.59 | A, C, D |
| ITGAM | ITGB2 | 0.75 | 0.71 | A, B, C, D |
| ITGB4 | ITGB6 | 0.75 | 0.28 | None |
| ITGA2 | ITGA4 | 0.74 | 0.46 | None |
| ITGA3 | ITGB8 | 0.72 | -0.002 | A, C, D |
| ITGAV | ITGA1 | 0.72 | 0.73 | None |
| ITGAV | ITGB1 | 0.71 | 0.76 | A, B, C, D |
| ITGA3 | ITGB4 | 0.70 | 0.19 | A, B, C, D |
| ITGA10 | ITGB4 | 0.69 | 0.17 | None |
| ITGA4 | ITGB7 | 0.68 | 0.42 | A, B, C, D |
| ITGA5 | ITGB3 | 0.68 | 0.60 | A, B, C, D |
| ITGA3 | ITGB6 | 0.66 | 0.27 | A, C, D |
| ITGA4 | ITGB8 | 0.64 | 0.26 | C, D |
| ITGA11 | ITGB8 | 0.64 | 0.09 | C, D |
| ITGA2 | ITGA11 | 0.64 | 0.44 | C, D |
| ITGA4 | ITGA11 | 0.62 | 0.45 | None |
| ITGA4 | ITGB6 | 0.62 | 0.21 | A, C, D |
| ITGA2 | ITGB7 | 0.62 | -0.06 | None |
| ITGA3 | ITGA4 | 0.61 | 0.12 | None |
| ITGA1 | ITGB1 | 0.61 | 0.73 | A, B, C, D |

*Red indicates a large difference in co-expression for the integrin pair (i.e., the difference between Pearson correlation R in the healthy and cancer samples was > 0.4)
**STRING-db associations (website checked on 11/21/23): A: coexpression; B: experimental/biochemical data; C: association in curated databases; D: co-mentioned in pubmed abstracts

**Supplementary Table 3:** Pearson correlation coefficients for integrin-integrin pairs that are highly correlated (Pearson R ≥ 0.6) in TCGA BRCA primary tumor samples, and their corresponding coefficients in GTEx*

| Integrin | Integrin | TCGA Primary Tumor R | GTEX R | STRING-db** Association? |
|---|---|---|---|---|
| ITGAX | ITGB2 | 0.85 | 0.28 | A, B, C, D |
| ITGA11 | ITGBL1 | 0.79 | 0.41 | A, D |
| ITGAV | ITGB1 | 0.76 | 0.71 | A, B, C, D |
| ITGA1 | ITGB1 | 0.73 | 0.61 | A, B, C, D |
| ITGAV | ITGA1 | 0.73 | 0.72 | None |
| ITGAM | ITGAX | 0.73 | 0.27 | None |
| ITGAM | ITGB2 | 0.71 | 0.75 | A, B, C, D |
| ITGAV | ITGB3 | 0.71 | 0.44 | A, B, C, D |
| ITGA1 | ITGA4 | 0.68 | 0.08 | None |
| ITGB2 | ITGB7 | 0.68 | 0.18 | None |
| ITGA1 | ITGB3 | 0.67 | 0.37 | A, C, D |
| ITGB1 | ITGB3 | 0.66 | 0.34 | None |
| ITGA1 | ITGA11 | 0.64 | -0.07 | A, C, D |
| ITGAV | ITGA4 | 0.65 | 0.18 | None |
| ITGA1 | ITGA8 | 0.65 | 0.35 | None |
| ITGA5 | ITGA11 | 0.63 | -0.05 | A, C, D |
| ITGA11 | ITGB1 | 0.62 | -0.16 | A, C, D |
| ITGAV | ITGA11 | 0.62 | 0.04 | None |
| ITGA4 | ITGB1 | 0.61 | -0.14 | A, B, C, D |
| ITGAV | ITGA2 | 0.60 | 0.04 | None |
| ITGAL | ITGAX | 0.61 | 0.57 | None |
| ITGA1 | ITGA5 | 0.61 | 0.50 | None |

*Red indicates a large difference in co-expression for the integrin pair (i.e., the difference between Pearson correlation R in the healthy and cancer samples was > 0.4)

**STRING-db associations (website checked on 11/21/23): A: coexpression; B: experimental/biochemical data; C: association in curated databases; D: co-mentioned in pubmed abstracts

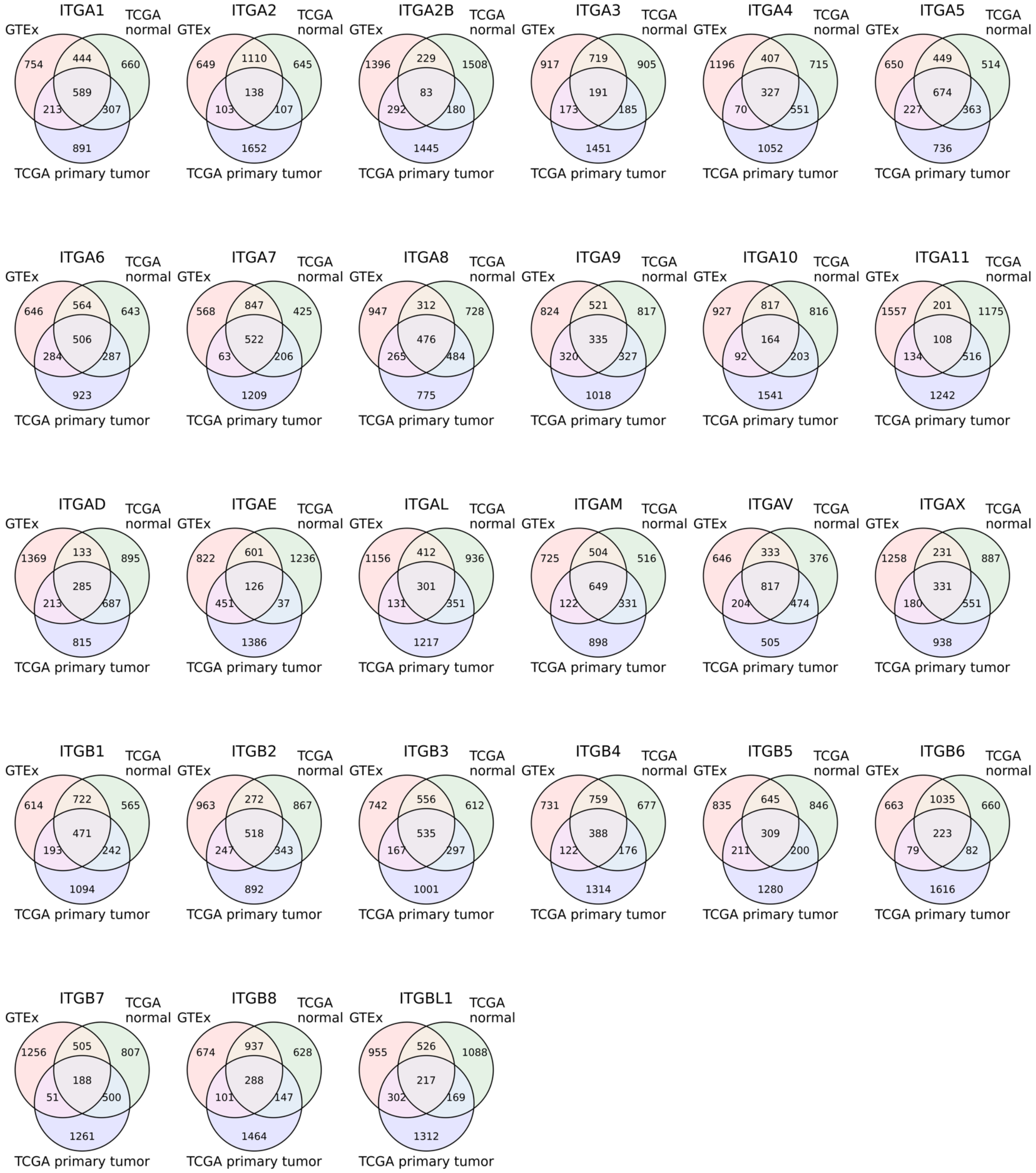


**Supplementary Fig. 11:** Venn diagrams of the overlap in genes that were co-expressed with each integrin in three datasets: GTEx breast, TCGA BRCA normal, TCGA BRCA primary tumor. For each integrin, the top 2000 protein-coding genes with the highest Pearson R were identified in each dataset, and the genes that overlapped in the three datasets were counted. For example, of the 2000 genes most co-expressed with ITGA1 in GTEx, 444 were also in the top 2000 genes for ITGA1 in TCGA primary tumor only, 213 were also in the top 2000 genes for ITGA1 in TCGA normal, and 589 were in the top 2000 genes in both TCGA datasets.

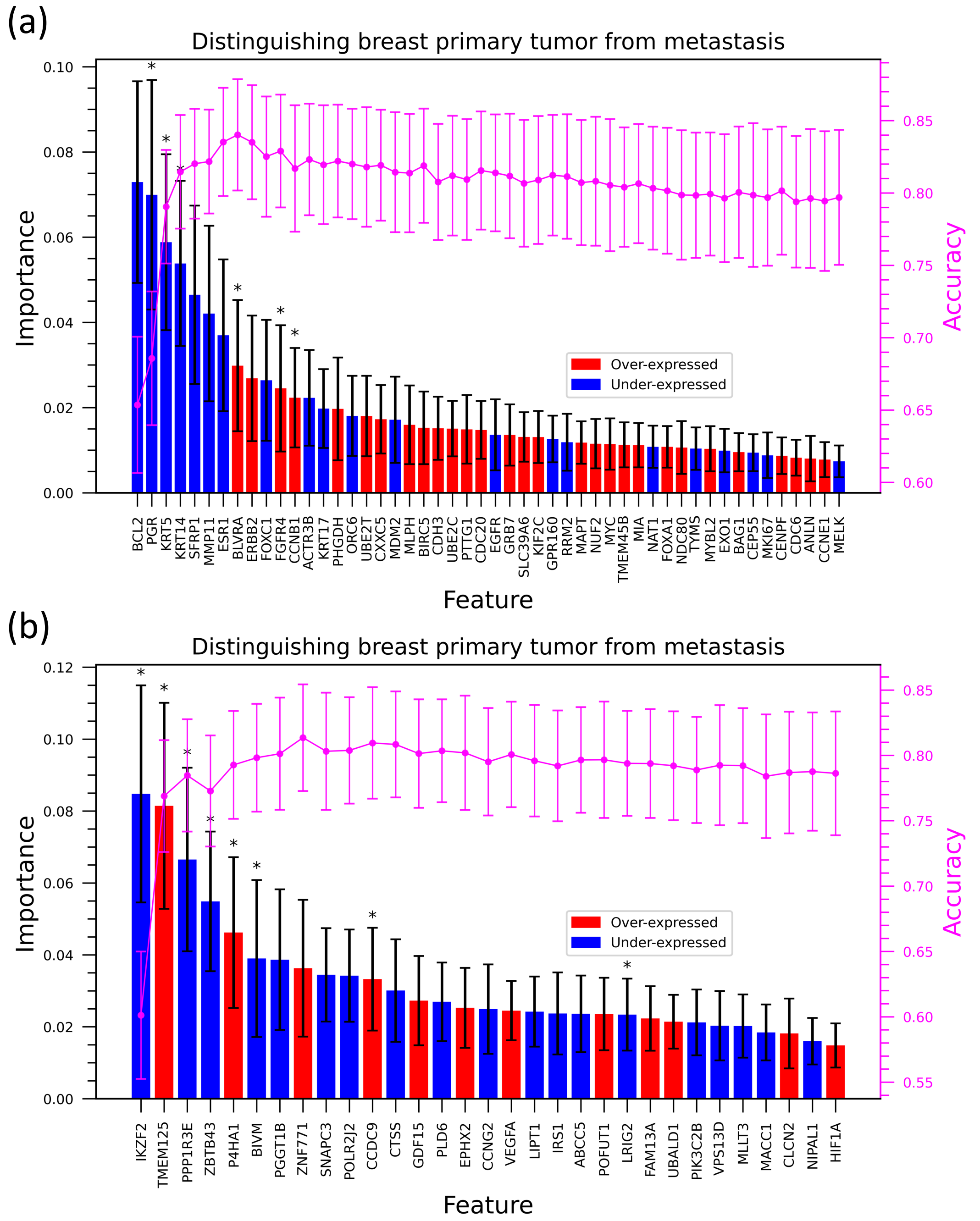


**Supplementary Fig. 12:** (a) Feature importance in a Random Forest model using a 50%/50% training/test split that classifies samples as primary breast tumor or metastatic tumor using the PAM50 gene set. The magenta line shows the change in the accuracy of the model as features are added one-by-one from the most important feature to the least. The accuracy of the model using all features is 0.80 ± 0.047 and AUROC is 0.86 ± 0.040. (b) Feature importance in a Random Forest model using a 50%/50% training/test split that classifies samples as primary breast tumor or metastatic tumor using the Donato gene set. The magenta line shows the change in the accuracy of the model as features are added one-by-one from the most important feature to the least. The accuracy of the model using all features is 0.78 ± 0.047 and AUROC is 0.82 ± 0.046.

**Supplementary Table 4:** Results of ANOVA test and Tukey post-hoc test for integrin expression in primary tumor test (breast) and 4 metastasis sites (liver, lymph node, brain, lung) in AURORA dataset (only integrins with significant expression differences are shown)

| Integrin | Bonferroni adjusted ANOVA p-values | Tukey post-hoc test* |
|---|---|---|
| **ITGA4** | 3E-7 | Brain & Breast (0.01) |
| | | Liver & Breast (0) |
| | | Liver & Lymph node (0.002) |
| **ITGA11** | 7E-7 | Brain & Breast (0.0004) |
| | | Liver & Breast (0) |
| | | Lung & Breast (0.05) |
| **ITGA7** | 1E-5 | Liver & Breast (0) |
| | | Liver & Brain (0) |
| | | Liver & Lymph node (0.002) |
| **ITGBL1** | 7E-5 | Brain & Breast (0.04) |
| | | Liver & Breast (0) |
| | | Lymph node & Breast (0.002) |
| **ITGAX** | 3E-4 | Brain & Breast (0.01) |
| | | Liver & Breast (0) |
| **ITGA8** | 6E-4 | Liver & Brain (0.0007) |
| | | Liver & Breast (0.0002) |
| | | Liver & Lung (0.0002) |
| | | Liver & Lymph node (0.04) |
| **ITGAM** | 9E-4 | Liver & Breast (0) |
| | | Liver & Lung (0.005) |
| | | Liver & Lymph node (0.009) |
| **ITGAE** | 8E-3 | Liver & Brain (0.01) |
| | | Liver & Breast (0.0004) |
| | | Liver & Lymph node (0.002) |
| **ITGAL** | 0.03 | Brain & Breast (0.03) |
| | | Liver & Breast (0.001) |
| **ITGB3** | 0.03 | Liver & Breast (0.002) |

*p-values are shown in parentheses

**Supplementary Table 5:** Summary of results in this work associated with each integrin

| Integrin | Healthy Tissue | Cancer Tissue | Healthy Breast/Breast Cancer | Breast Cancer Metastasis |
|---|---|---|---|---|
| **ITGA1** | | 2nd ranked feature in LUSC-vs-all classifier | Conserved co-expression with ITGAV and ITGB1; higher co-expression with integrins in cancer | |
| **ITGA2** | | | Conserved co-expression with ITGAV; higher co-expression with integrins in healthy samples | |
| **ITGA2B** | 1st feature in testis-vs-all classifier | 3rd feature in multiclass classifier; 1st feature in TGCT-vs-all classifier | Not co-expressed with any other integrin in healthy or cancer samples. | |
| **ITGA3** | | 1st feature in multiclass and BRCA-vs-all classifiers | Co-expressed with several integrins in healthy samples only | |
| **ITGA4** | | | Co-expressed with several integrins in healthy and cancer samples | Significantly lower expression in metastatic breast cancer than primary breast cancer |
| **ITGA5** | | Low variation in mean expression across cancer tissues. | Co-expressed with ITGA3 in healthy and with ITGA1 and ITGA11 in cancer | |
| **ITGA6** | 1st feature in liver-vs-all and breast-vs-all classifiers | 1st feature in COAD-vs-all classifier | Not co-expressed with any other integrin in healthy or cancer samples | |
| **ITGA7** | | | 1st feature in distinguishing healthy vs breast samples; remains 1st feature using other gene sets in model | |
| **ITGA8** | 2nd feature in multiclass, 1st feature in prostate-vs-all and stomach-vs-all classifiers | 1st feature in PRAD-vs-all classifier | | |
| **ITGA9** | 1st feature in pancreas-vs-all classifier | | Not co-expressed with any other integrin in healthy or cancer samples | |
| **ITGA10** | | | Co-expressed with ITGB4 in healthy samples | Least difference in expression between |

| | | | | |
|---|---|---|---|---|
| | | | | primary and metastatic breast cancer |
| **ITGA11** | | | Co-expressed with several integrins in healthy and cancer samples, but co-expression is not conserved | Significantly lower expression in metastatic breast cancer than primary breast cancer |
| **ITGAD** | Low expression across tissues | Low expression across cancers | Not co-expressed with any other integrin in healthy or cancer samples | Significantly lower expression in metastatic breast cancer than primary breast cancer |
| **ITGAE** | | | Not co-expressed with any other integrin in healthy or cancer samples | |
| **ITGAL** | | | | Significantly lower expression in metastatic breast cancer than primary breast cancer |
| **ITGAM** | | | Conserved co-expression with ITGB2 | |
| **ITGAV** | | | Conserved co-expression with ITGA1 and ITGB1; higher co-expression with integrins in cancer | |
| **ITGAX** | 1st feature in lung-vs-all classifier | | Co-expressed with other leukocyte integrins in cancer | Significantly lower expression in metastatic breast cancer than primary breast cancer |
| **ITGB1** | High mean expression in all tissues, and low variation in expression across tissues | High mean expression in all cancer, and low variation in expression across tissues | Conserved co-expression with ITGA1 and ITGAV, also co-expressed with other integrins including ITGA4 and ITGA11 in cancer | Highest mean expression in primary breast tumor and breast metastasis |
| **ITGB2** | | | Conserved co-expression with ITGAM | |
| **ITGB3** | | Relatively low variation in mean expression across cancers | | |
| **ITGB4** | 1st feature in multiclass classifier | 1st feature in STAD-vs-all classifer | Co-expressed with several integrins in healthy samples, but not co-expressed with any integrin in cancer samples | |

| | | | | |
|---|---|---|---|---|
| **ITGB5** | | High mean expression in all cancers but low variation in mean expression across cancers | Not co-expressed with any other integrin in healthy or cancer samples | |
| **ITGB6** | 3rd feature in multiclass classifier | 2nd feature in multiclass and 1st feature in LIHC-vs-all and PAAD-vs-all classifiers | Co-expressed with several integrins in healthy samples only | |
| **ITGB7** | | | Co-expressed with ITGA2, ITGA4, and ITGAL in healthy samples and with ITGB2 in cancer samples | |
| **ITGB8** | | 1st feature in LUSC-vs-all classifier | Co-expressed with several integrins in healthy samples only | |
| **ITGBL1** | 1st feature in colon-vs-all classifier | | | Significantly lower expression in metastatic breast cancer than primary breast cancer |
